# USH2A is a skin end-organ protein necessary for vibration sensing in mice and humans

**DOI:** 10.1101/2020.07.01.180919

**Authors:** Fred Schwaller, Valérie Bégay, Gema García-García, Francisco J. Taberner, Rabih Moshourab, Brennan McDonald, Trevor Docter, Johannes Kühnemund, Julia Ojeda-Alonso, Ricardo Paricio-Montesinos, Stefan G. Lechner, James F.A. Poulet, Jose M. Millan, Gary R. Lewin

## Abstract

Fingertip mechanoreceptors comprise sensory neuron endings together with specialized skin cells that form the end-organ. Exquisitely sensitive vibration-sensing neurons are associated with Meissner’s corpuscles and Pacinian corpuscles^1^. Such end-organ structures have been recognized for more than 160 years, but their exact functions have remained a matter of speculation. Here we examined the role of USH2A in touch sensation in humans and mice. The *USH2A* gene encodes a transmembrane protein with a very large extracellular domain. Pathogenic *USH2A* mutations cause Usher syndrome associated with hearing loss and visual impairment^2^. We show that patients with biallelic pathogenic *USH2A* mutations also have profound impairments in vibrotactile touch perception. Similarly, mice lacking the USH2A protein showed severe deficits in a forepaw vibrotactile discrimination task. Forepaw rapidly-adapting mechanoreceptors (RAMs) recorded from *Ush2a*^*−/−*^ mice innervating Meissner’s corpuscles showed profound reductions in their vibration sensitivity. However, the USH2A protein was not expressed in sensory neurons, but was found in specialized terminal Schwann cells in Meissner’s corpuscles. Loss of this large extracellular tether-like protein in corpuscular end-organs innervated by RAMs was sufficient to reduce the vibration sensitivity of mechanoreceptors. Thus, USH2A expressed in corpuscular end-organs associated with vibration sensing is required to properly perceive vibration. We propose that cells within the corpuscle present a tether-like protein that may link to mechanosensitive channels in sensory endings to facilitate small amplitude vibration detection essential for the perception of fine textured surfaces.

---

There are several lines of evidence showing common molecular and genetic factors shared by the senses of hearing and touch^3–6^. For example, some patients with Usher syndrome, which is characterized by severe to moderate hearing impairment followed by late onset visual impairment, also show deficits in touch perception^3^. Here we decided to focus on the *USH2A* gene because mutations in this gene are one of the most frequent causes of Usher syndrome^7^. The USH2A protein is a large transmembrane protein that has been proposed to form the tether-like ankle link at the base of hair cell stereocilila^8,9^. We recruited 13 patients with bi-allelic truncation or null mutations in *USH2A* gene and asked whether such patients display specific touch deficits compared to a cohort of 65 age and gender matched healthy controls (Supplementary Table 1; Fig. 1a). We used quantitative sensory testing including a two interval forced choice procedure to measure vibration detection threshold (VDT) on the little finger^4^. *USH2A* patients had robustly elevated perceptual detection thresholds for both 10Hz and 125Hz vibration stimuli (Fig 1b-c, Supplementary Table 2). Mean vibration detection thresholds for 125Hz were >4 times higher in the patient cohort, and 7/13 patients had Z-scores below −1.96 (~2 standard deviations different from the control mean; Fig 1b, c; Supplementary Table 2). Detection of 10Hz stimuli was also significantly poorer (>1.5 times) in the patient group compared to controls (Fig 1b, c). The patient cohort displayed significantly elevated cool detection thresholds (CDT), mean 1.6 ^°^C compared to 1.0 ^°^C in the control cohort (Fig 1d, Supplementary Tables 2, 3). Changes in thermal perception thresholds have previously been observed in hearing- and touch-impaired patients, many of whom do not have pathogenic *USH2A* mutations^3,4^. Importantly, the *USH2A-*patients did not substantially differ with respect to the control population in punctate stimulus detection thresholds, pinprick pain detection thresholds, and heat and cold pain thresholds (Fig 1b, Supplementary Table 2). *USH2A*-patients showed slightly better performance in the tactile acuity test compared to the control cohort (Fig 1b, Extended data Table 2). Amongst the main traits that showed significant loss of function (VDT 10 HZ, VDT 125Hz and CDT) there was a wide range of severities that may be loosely related to the genotype (Supplementary Table 3). Thus, *USH2A* loss-of-function mutations in humans are strongly associated with a selective impairment of vibration perception.

**Fig. 1.**
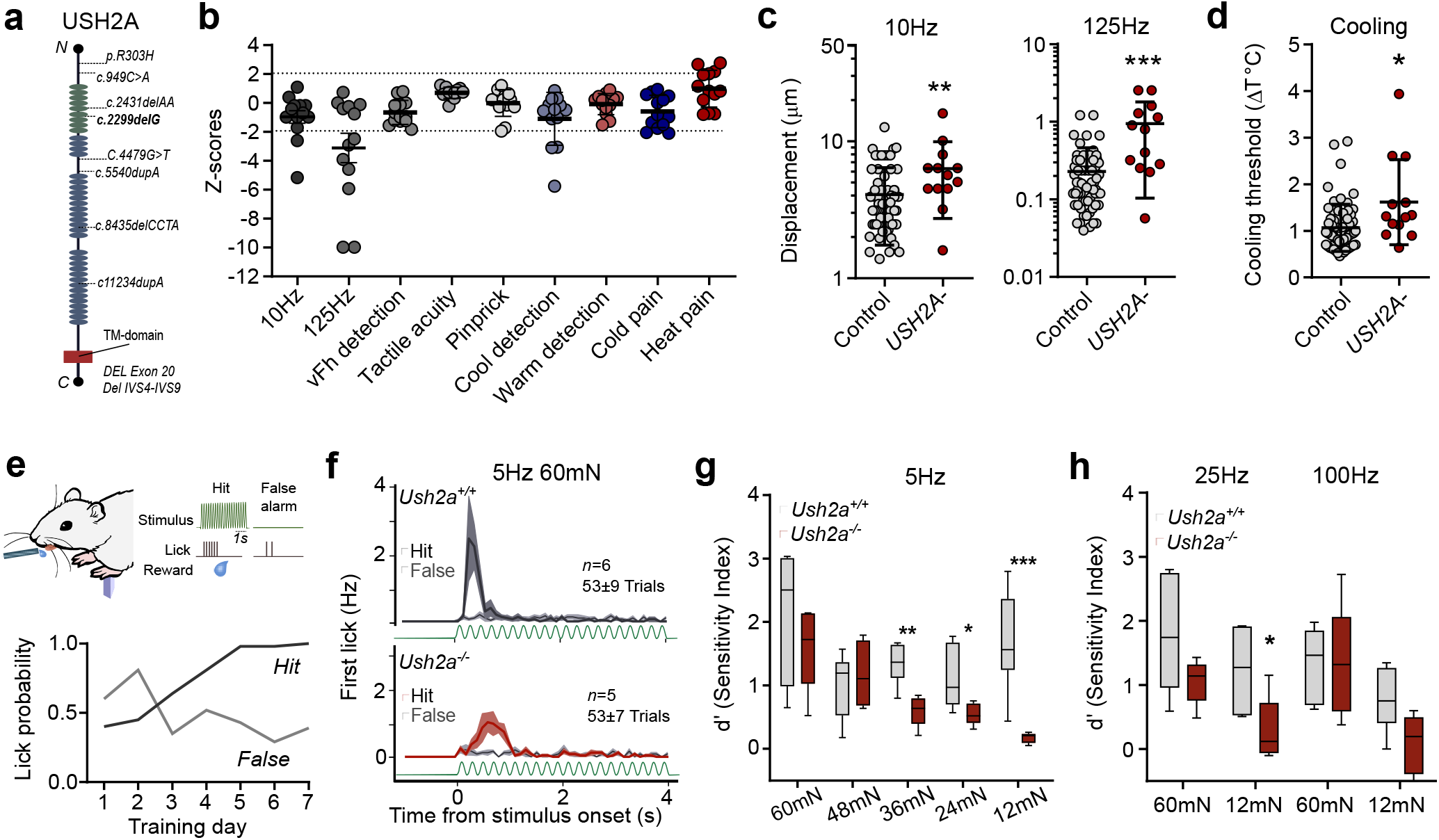
*USH2A/Ush2a* mutations impair cutaneous vibration perception in humans and mice. **a,** Structure of USH2A protein with locations of truncation and loss of function (LoF) gene mutations present in the cohort of 13 patients with Usher Syndrome type II. **b,** Psychophysical thresholds of patients with *USH2A* mutations normalized as Z-scores. Data lines indicate upper and lower boundaries of 95% confidence interval of the normal distribution of healthy subjects. **c,**10Hz (left) vibration perceptual thresholds were higher in patients with *USH2A* mutations *(n* = 13) compared to healthy controls (*n* = 65) (\*\**P* <0.005 Mann-Whitney U-test). 125Hz (right) vibration perception thresholds were also higher in patients with *USH2A* mutations (\*\*\**P* <0.001 Mann-Whitney U-test). Data = mean± S.D. **d,** Cooling detection thresholds were higher in patients compared to control (\**P* <0.01 Mann Whitney U-test). Data = mean± S.D. **e,** Mouse vibration perceptual task experimental set up (upper panel) and example hit and false alarm rates (lower panel) from one *Ush2a*^*+/+*^ mouse during the forepaw 60mN 5Hz vibration stimulus learning phase. **f,** Latency histograms of first lick responses from vibration-trained *Ush2a*^*+/+*^ (top; *n* = 6) and *Ush2a*^*−/−*^ mice (bottom; *n* = 5) on training day 7. Data = mean± S.D. **g,** Perceptual sensitivity index (d’) measurements revealed that *Ush2a*^*−/−*^ mice had poorer performance at low-amplitude 5Hz vibrations than *Ush2a*^*+/+*^ mice (*,**,***P <0.05, 0.01, 0.001, two-way ANOVA with Bonferroni post-hoc analysis. **h,** *Ush2a*^*−/−*^ mice had poorer performance at low-amplitude 25Hz vibrations (*P <0.05 two-way ANOVA with Bonferroni post-hoc analysis).

We next asked whether genetic ablation of *Ush2a* in mice impairs forepaw vibration detection in behaving animals. We adapted a goal-directed sensory perception task^10,11^ and trained water-restricted and head restrained *Ush2a*^*+/+*^ and *Ush2a*^*−/−*^ mice^12^ to report a 5Hz forepaw vibration stimulus for a water reward^13^ (Fig 1e). Mice learned to report a 60mN 5Hz forepaw vibration over a 7-day training period (Fig 1e, f, Extended data Fig. 1a-c). Decreasing stimulus amplitudes revealed that control mice could detect 5Hz vibration with amplitudes greater than 12mN, but *Ush2a*^*−/−*^ mice were only able to detect the same stimulus at amplitudes greater than 24mN (Fig 1g, Extended data Fig. 1d), indicating that *Ush2a*^*−/−*^ mice had significantly higher 5Hz vibration perceptual thresholds than control mice. Similarly, *Ush2a*^*−/−*^ mice could not detect low amplitude 12mN 25Hz vibration stimuli that were reliably reported by control mice (Fig 1g,h, Extended data Fig 1g). To compare directly the performance of mice in the 5Hz vibration detection task we calculated the sensitivity index (d’, see methods) and found that wild type mice performed significantly better than *Ush2a*^*−/−*^ mice when stimuli were applied with amplitudes ranging from 36 to 6mN (Fig 1h). The sensitivity index was also significantly different for the 25 Hz stimulus, but was not significant for the 100 Hz stimulus (Fig 1g,h; Extended data Fig 1 g,h). Since *Ush2a*^*−/−*^ mice have hearing impairments^12^, we tested whether the mice lick in response to any noise made by the mechanical stimulator. By placing the mechanical stimulator a short distance away from the forepaw at the end of the test session, we showed that both control and *Ush2a*^*−/−*^ mice did not respond to sound of the stimulator with licking greater than the false lick rate (Extended data Fig 1i). Consistent with previous studies *Ush2a*^*−/−*^ mice also showed hearing impairments in a paired pulse inhibition hearing test (Extended data Fig 2). Thus, the absence of the USH2A protein in mice and in humans is associated with profound deficits in vibration perception and mild hearing loss.

USH2A is a large extracellular tether-like protein that is expressed during development in the inner ear at the base of hair cell stereocilia, but does not appear to contribute to mechanotransduction at the tip link of stereocilia^7–9,12,14^. Mechanotransduction channels essential for touch are primarily located in sensory neurons^15,16^, however, the USH2A protein was undetectable in adult dorsal root ganglia (DRG) neurons (Extended data Fig. 3a), confirmed by published RNA sequencing datasets^17^. Using an RNAscope probe specific for Ush2a mRNA we found no signal in the DRG, the same probe did label Meissner corpuscles in the skin (Extended data Fig 3a). We thus hypothesised that USH2A may be expressed in skin cells associated with vibration sensitive sensory neurons. Indeed, we detected USH2A immunofluorescence in adult forepaw glabrous skin that was restricted to the capsule of the Meissner corpuscle (Fig 2a,b,c Extended data Fig. 3b,c). The USH2A immunofluorescence signal did not, however, overlap with Neurofilament-200 (NF200, a marker of myelinated sensory axons) in the dermis or hypodermis (Fig 2a, b, Extended data Fig. 3b, d). Instead, USH2A immunofluorescence was co-localized with labelling for S100 (a marker of terminal Schwann cells) surrounding Aβ-fiber neurites in the Meissner corpuscle capsule. However, outside the Meissner corpuscle we could not detect a signal for USH2A in myelinating Schwann cells proximal to the end-organ, a cell type recently reported to be mechanosensitive^18^ (Fig 2a, b, Extended data Fig. 3c). USH2A immunofluorescence was absent from Meissner’s corpuscles in *Ush2a*^*−/−*^ mice (Extended data Fig. 3d). Interestingly, high magnification images revealed USH2A immunofluorescence signals in the terminal Schwann cell in close proximity to the sensory ending (Fig 1c arrows, see also Z-stack animations of further examples in Supplementary Movie 1-3). Together these data demonstrate that the USH2A protein is present in adult skin sensory end-organs where it plays a role in vibration-sensing.

**Fig. 2.**
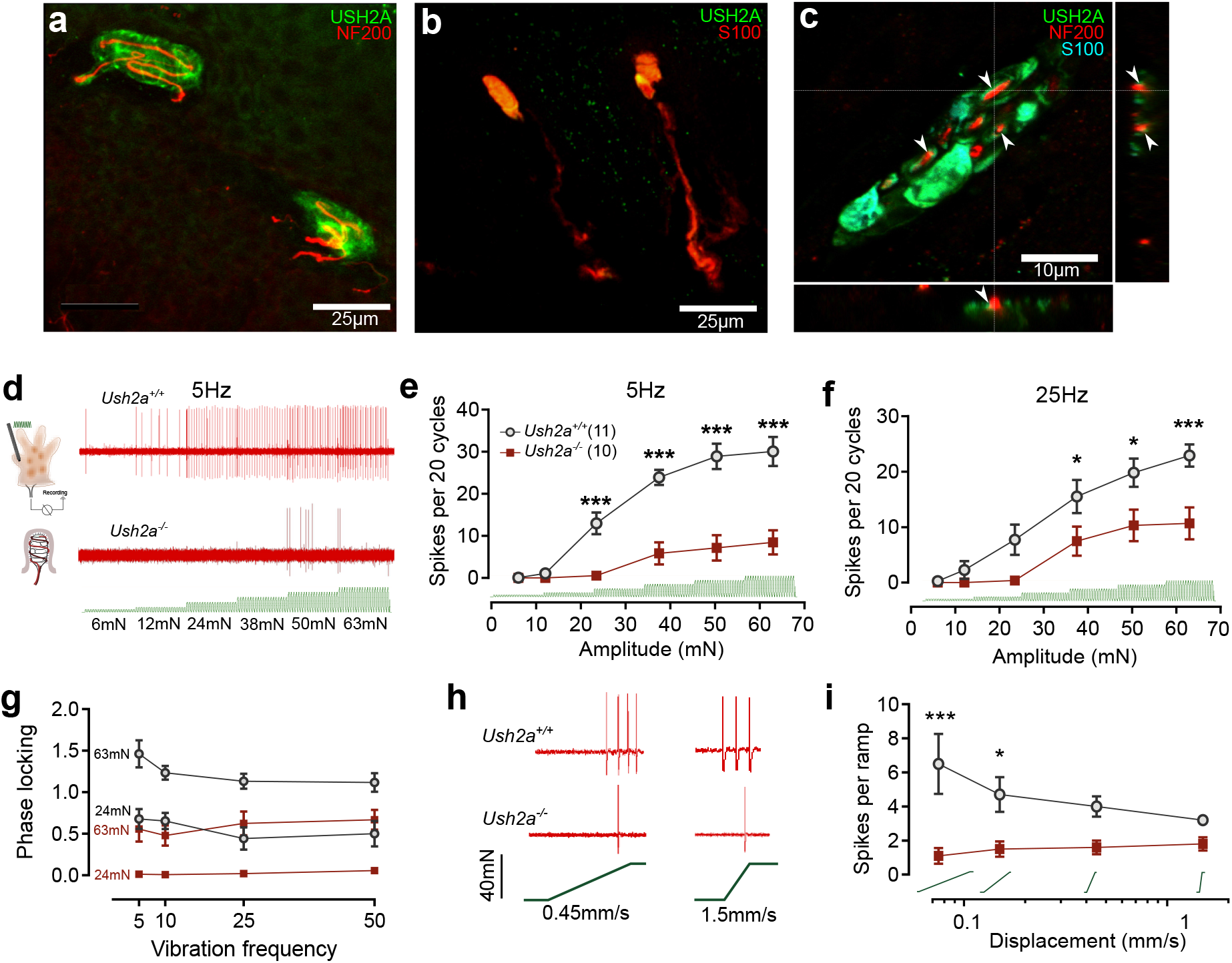
USH2A is expressed in Meissner corpuscles and facilitates their vibrosensitivity. **a,** USH2A immunolabeling (green) was present in Meissner corpuscles at the dermal and epidermal boundary sections of mouse forepaw glabrous skin, but did not colocalise with NF200 (red). **b,** USH2A (green) colocalised with S100 (red) in the bulb of Meissner corpuscles (arrowhead, inset with USH2A channel only). **c,** 63x magnification Z-stack of an USH2A+ Meissner corpuscle. USH2A overlapped with S100 in terminal schwann cells and appeared to wrap around NF200+ nerve endings (arrows). **d,** RAM Aβ-fibers innervating Meissner corpuscles recorded using the *ex vivo* forepaw glabrous skin nerve preparation. Example traces of single RAMs in *Ush2a*^*+/+*^ (top) and *Ush2a*^*−/−*^ (bottom) mice in response to 5Hz vibration. **e,** RAMs in *Ush2a*^*−/−*^ mice (*n* = 10 afferents from 4 mice) had significantly reduced firing activity to 5Hz vibration compared to *Ush2a*^*+/+*^ mice (*n* = 11 afferents from 5 mice; ***P < 0.001, two-way ANOVA with Bonferroni post-hoc analysis). **f**, 25Hz vibration sensitivity was also impaired in *Ush2a*^*−/−*^ RAMs (**P < 0.001, two-way ANOVA with Bonferroni post-hoc analysis)**. g,** Phase locking (number of spikes per sinusoidal wave) was impaired in *Ush2a*^*−/−*^ mice at two different amplitudes of 5Hz vibration: 63mN and 24mN. Data = mean± S.E.M. **h**, Example ramp responses of individual RAMs from *Ush2a*^*+/+*^ and *Ush2a*^*−/−*^ mice. **i,** RAM responses to ramp stimuli were impaired in *Ush2a*^*−/−*^ mice (*P < 0.05, two-way ANOVA with Bonferroni post-hoc analysis). Data = mean± S.E.M.

Glabrous skin Meissner corpuscles are innervated by Aβ-fiber RAMs that are tuned to detect small amplitude skin vibration below 80Hz^1,5,19^. We next recorded extracellular spike activity from single mechanoreceptors in *Ush2a*^*−/−*^ mice using *ex vivo* skin nerve preparations^20^. Meissner corpuscle-associated RAMs (only found in glabrous skin) had severely impaired vibration sensitivity in *Ush2a*^*−/−*^ mice compared to *Ush2a*^*+/+*^ mice (Fig 2d-g, Extended data Figs. 4a-e). Notably, forepaw Meissner corpuscle RAMs in *Ush2a*^*−/−*^ mice were significantly less responsive to 5Hz, 10Hz, 25Hz and 50Hz vibration stimuli. Control RAMs show phase-locked spiking to each sinusoid at low forces, and often fired more than one spike per sinusoid at higher forces (Fig 2 e-g.). In contrast, RAMs in *Ush2a*^*−/−*^ mice showed incomplete phase locking even at high forces at all frequencies tested (Fig. 2g, Extended data Figs. 4a-e). Using ramp stimuli of different velocities^20,21^ we also observed that RAMs from *Ush2a*^*−/−*^ mice failed to fire spikes with slow ramp velocities compared to RAMs from controls (Fig. 2h,I, Extended data Fig. 4h). As we previously showed, forepaw RAMs were more mechanosensitive compared to hindpaw RAMs^20^ (Extended data Fig 4i,j). However, even though RAMs in glabrous hind paw skin were not as vibration sensitive as those in the forepaw they showed clear and significant deficits in vibration sensitivity in *Ush2a*^*−/−*^ mice (Extended Fig 4f,g). Thus, the presence of the USH2A protein in terminal Schwann cells of the Meissner corpuscle appears to be necessary for normal vibration sensitivity.

We next asked if the USH2A protein could play a role in regulating the sensitivity of other RAMs outside the glabrous skin^22^. In the hairy hind paw skin, USH2A immunofluorescence localised around most hair follicles with lanceolate endings and in some, but not all, hair follicles with circumferential endings (Fig 3a-d, Extended data Fig. 5a-d). In more than 80% of hair follicles with lanceolate endingsUSH2A immunofluorescence co-localised with S100 in the terminal Schwann cells, which surround the NF200+ lanceolate endings (Fig 3a, Extended data Fig. 5a). We found USH2A immunofluorescence was present in terminal Schwann cells associated with around 50% of identified Guard hairs and only 43% of follicles with circumferential endings (Fig 3d). USH2A immunofluorescence was localized at sites where tether-like proteins have been visualized^23^, again published single cell sequencing data indicates that *Ush2a* mRNA is present in cells surrounding the hair follicle (http://linnarssonlab.org/epidermis/)^24^. We tested the mechanosensitivity of Aβ-fiber RAMs innervating hair follicles in the hind paw by recording their extracellular spike activity using the hind paw hairy skin nerve preparation. On average, hair follicle Aβ-fiber RAMs in *Ush2a*^*−/−*^ mice had impaired vibration sensitivity to 5Hz, but not to 25Hz or 50Hz sinusoids, compared to controls (Fig 3 e-g, Extended data Fig 6a-c). The mild and selective deficit in mechanosensitivity to very low frequency vibration suggests a relatively minor role for the USH2A protein in these hair follicle associated mechanoreceptors.

**Fig. 3.**
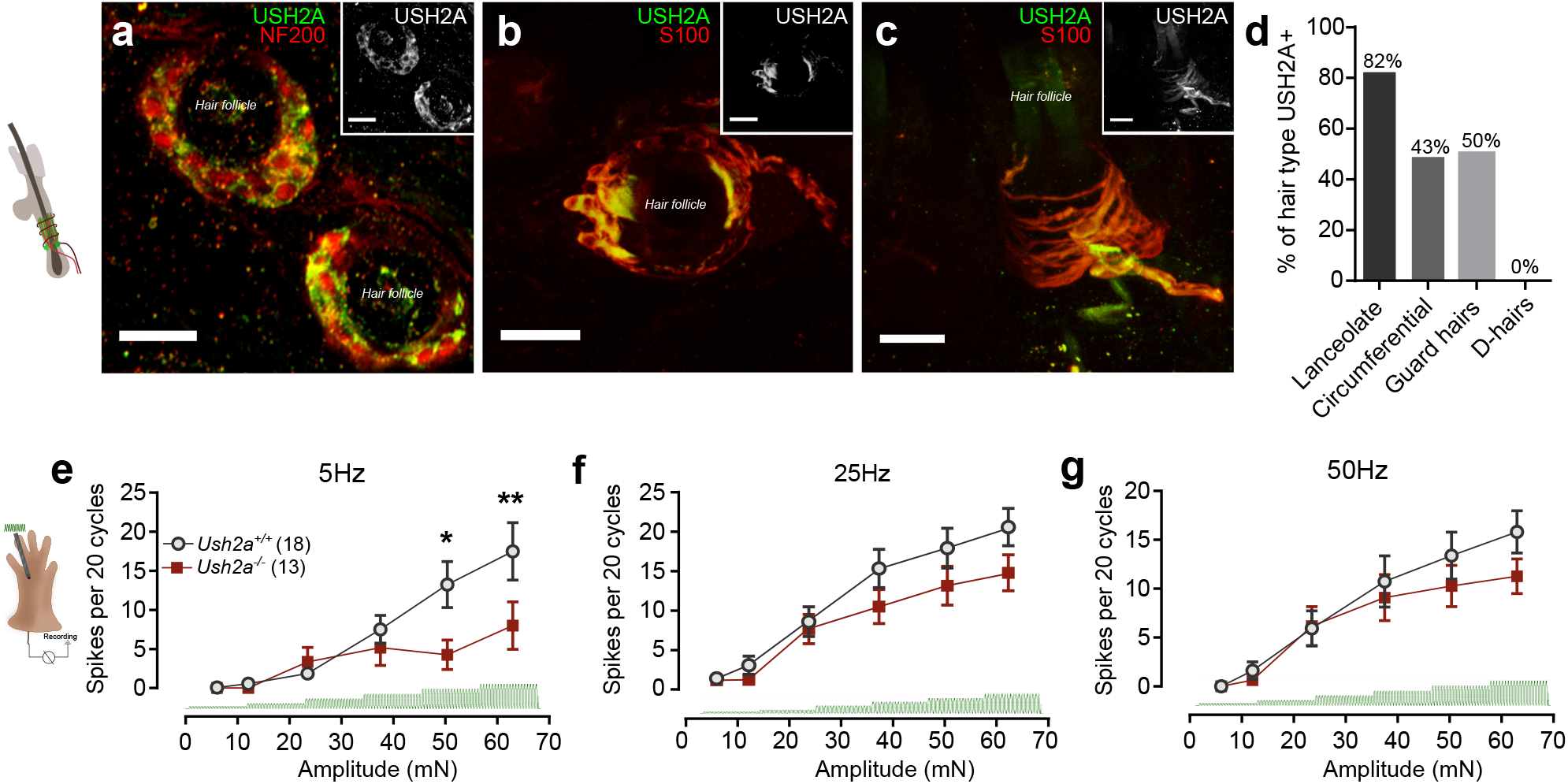
USH2A is expressed in some hair follicles, and facilitates their vibrosensitivity. **a,** USH2A (green) in hair follicle end organs in oblique cross sections of hindpaw hairy skin surrounding NF200+ lanceolate nerve endings. **b**, USH2A (green) colocalised with S100 (red) in terminal Schwann cells surrounding lanceolate endings in hair follicles in wholemounted hindpaw hairy skin. **c,** USH2A also colocalised with S100 (red) around some circumferential endings. **d,** Quantification of USH2A expression in different types of hair follicle endings. **e,** RAMs innervating hair follicles recorded using the *ex vivo* hindpaw hairy skin nerve preparation in *Ush2a*^*−/−*^ mice (*n* = 13 neurons from 4 mice) had significantly reduced 5Hz vibration-evoked firing activity compared to *Ush2a*^*+/+*^ mice (*n* = 18 neurons from 5 mice). **f,**25Hz and **g,**50Hz vibration sensitivity was not significantly different between *Ush2a*^*+/+*^ and *Ush2a*^*−/−*^ mice. Data = mean± SEM. All scale bars 25μm.

For the mouse behavior we observed little or no deficit with a 100Hz (Fig 1h), but nevertheless wished to test if other vibration sensing neurons show deficits in the absence of USH2A. Pacinian corpuscle mechanoreceptors are high frequency vibration sensors^1^. In humans Pacinian corpuscle mechanoreceptors are termed type II RAMs and are common in the hand where they respond best to vibration frequencies above 300 Hz^25^. There are few studies on murine Pacinian corpuscle mechanoreceptors, but these receptors are predominantly found on the interosseous membrane along the long bones in mice^26,27,28^. Although mice appear to perceive high frequency vibration^28^, there have so far been no published recordings from Pacinian corpuscle mechanoreceptors in this species. Here we used a newly established mouse ex vivo preparation to record from mechanoreceptors innervating Pacinian corpuscles located around the hindlimb fibula bone. The vibration sensitivity of mouse Pacinian mechanoreceptors was probed using a series of sinusoidal stimulus trains with linearly increasing amplitudes. Using these stimuli we calculated the threshold amplitude for each frequency in a range from 40-480Hz and found that, as in other species^29,30^, these RAMs are preferentially tuned to frequencies > 300 Hz (Fig 4a,b). We could also record from Pacinian mechanoreceptors in *Ush2a*^*−/−*^ and *Ush2a*^*+/+*^ mice and found that there appeared to be no effect of the mutation on the sensitivity of these receptors to vibration at any frequency tested Thus, the tuning curves from *Ush2a*^*−/−*^ mice (on a CBA/CaJ background) did not differ from those recorded in *Ush2a*^*+/+*^ littermate controls (Fig 4 a,b). Furthermore, tuning curves recorded from Pacinian mechanoreceptors taken from C57Bl6/N mice also did not differ from those found in the *Ush2a*^*−/−*^ mice. Thus, the absence of the USH2A protein from mouse Pacinian corpuscle does not appear to lead to a decrease in the sensitivity of these mechanoreceptors to vibration.

**Fig. 4.**
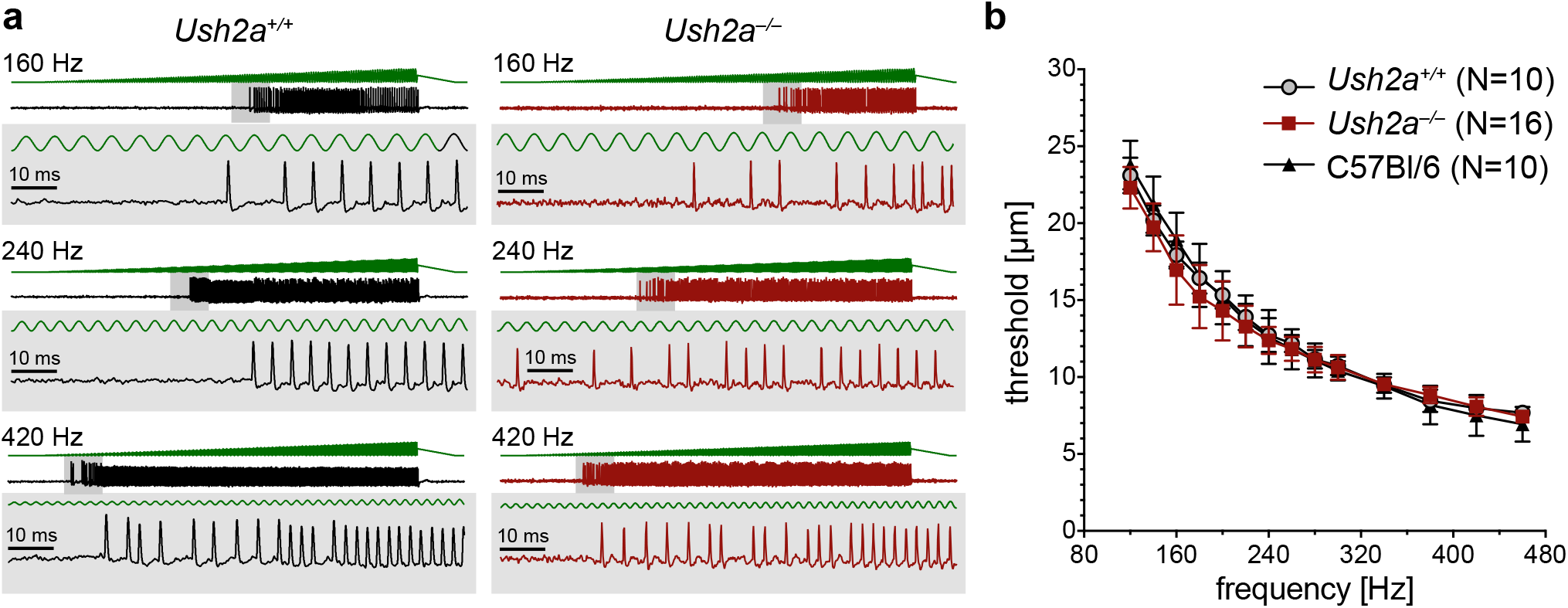
Pacinian corpuscle vibration sensitivity is unchanged in Ush2a−/− mice. **a**, Pacinian afferents were stimulated by applying a series of sinusoidal mechanical stimuli (duration 1s; linearly increasing amplitude *d*Amp/*d*t = 30μm/s; maximum amplitude 30μm) with increasing frequencies (40-480 Hz) to the distal end of the fibula. Examples of responses evoked by 160 Hz, 240 Hz and 420 Hz stimulation of Pacinian afferents from Ush2a^−/−^ mice and Ush2a^+/+^ littermate controls are shown in the right and left panel, respectively. The insets show magnified views of the grey shaded areas. **b**, Frequency tuning curves (mechanical activation threshold vs. stimulation frequency) of Pacinian afferents from *Ush2a*^*+/+*^ (black squares), *Ush2a^−/−^* (red circles) and C57Bl/6N (grey triangles) mice. Data = mean± SEM

Other cutaneous end organs such as Aβ-fiber slowly adapting mechanoreceptor (SAM)-Merkel cell complexes and Aδ-fiber D-hairs are also vibration sensitive. We next tested whether USH2A is present in these structures and contributes to their mechanosensitivity. USH2A immunofluorescence was not detected in CK20+ Merkel cell complexes surrounding guard hairs in the back skin (Fig 5a, Extended data Fig 5d) nor around the endings of D-hair receptors that innervate a group of fine hairs in the centre of the glabrous hind paw skin (Fig 3d, Fig 5c, Extended data Fig 5e)^20^. Using the *ex vivo* skin nerve recordings we found that the mechanosensitivity of SAMs (Fig 5b, Extended data Fig 6a-e) and D-hairs were not impaired in *Ush2a*^*−/−*^ mice (Fig 5d, Extended data Fig 6f-i). Additionally, the mechanosensitivity of SAMs and Aδ- and C-fiber nociceptors to high amplitude ramp and hold stimuli were also not impaired in *Ush2a*^*−/−*^ mice (Fig 5e-g). Consistent with these electrophysiological findings *Ush2a*^*−/−*^ mice displayed comparable behavioral forepaw brush-evoked responses and nocifensive withdrawal thresholds to control *Ush2a*^*+/+*^ mice (Fig 5h,i). These data from the mouse are thus in good agreement with studies with human patients indicating that USH2A loss of function alleles are associated specifically with vibration perception and not with other somatosensory modalities including pain (Fig 1). However, *USH2A*-patients did show mild but significant deficits in cool detection (Fig 1b,d). We thus asked if the sensory coding of thermal stimuli was altered in the absence of USH2A. In mice, skin warming and cooling are signalled by polymodal C-fibers that respond to small increases or decreases in temperature^10,11^. We recorded from polymodal C-fibers and classified them according to their response to mechanical, warm and cooling stimuli. We found the same proportion of C-mechanoheat, C-mechanocold, C-mechanoheatcold and C-cold primary afferents in *Ush2a*^*−/−*^ mice compared to their littermate controls (Extended data Fig 7a). Moreover, we used slow warming and cooling ramps^11^ (1°C/sec) to quantify thermosensitivity and found no deficits in cool or warm driven activity in mice lacking USH2A (Extended data Fig 7b,c). These data suggest that deficits in thermal perception associated with loss of function mutations in the *USH2A* gene are not due to functional changes in thermosensory afferents.

**Fig. 5.**
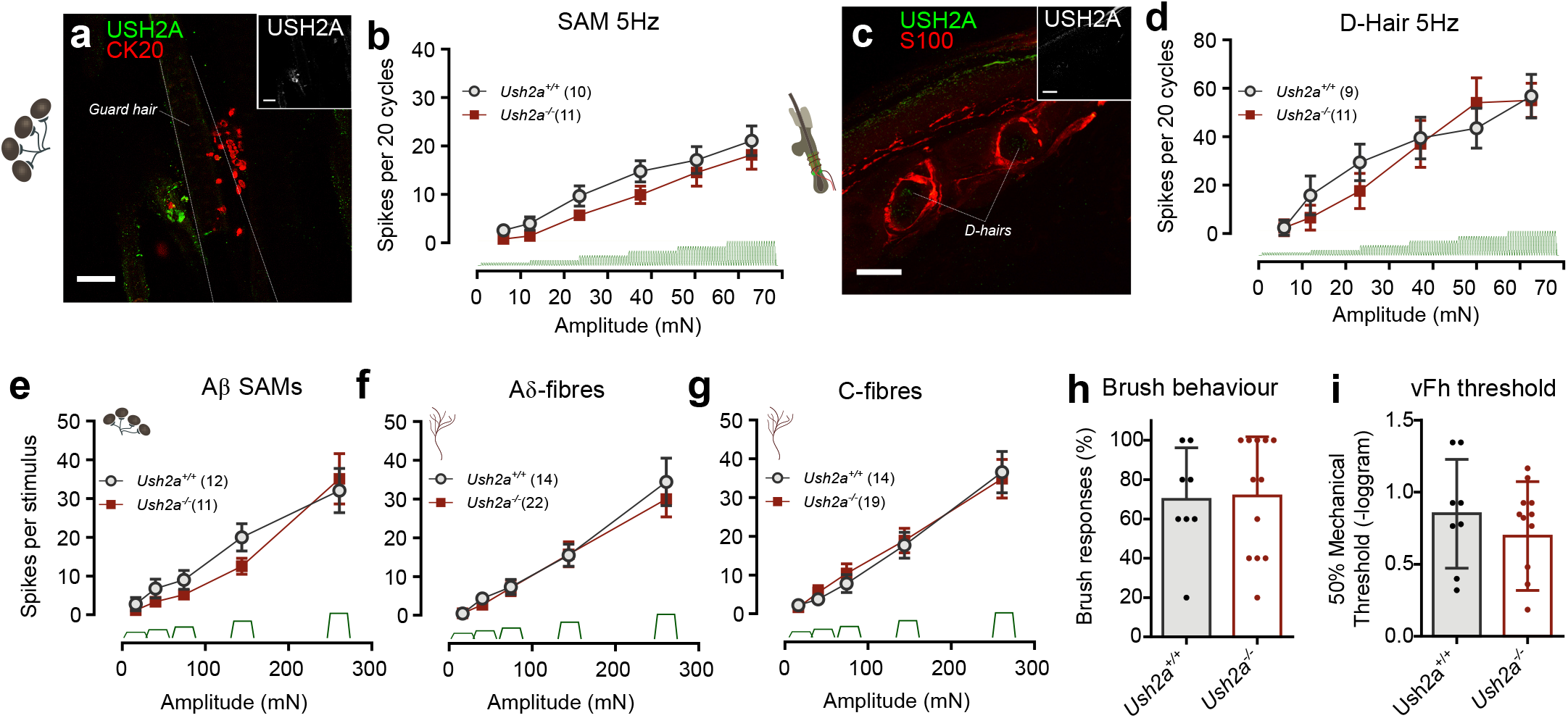
D-hair and SAM end organs do not express USH2A, and have intact mechanosensitivity in *Ush2a*^*−/−*^ mice. **a,** USH2A immunolabelling (green) does not colocalise with CK20 (red) in guard hair Merkel cell complexes in the mouse hairy back skin. **b**, Slowly adapting mechanoreceptive (SAM) Aβ-fibers that innervate Merkel cells in the glabrous forepaw skin of *Ush2a*^*−/−*^ mice (*n* = 11 neurons from 4 mice) show comparable 5Hz vibration sensitivity with *Ush2A+/+* mice (*n* = 10 neurons from 5 mice; *P* > 0.05, two-way ANOVA with Bonferroni post-hoc analysis). **c,** USH2A is absent from Aδ-fiber D-hairs in the glabrous hindpaw skin. **d**, Aδ-fiber D-hair 5Hz vibration sensitivity is not significantly different between *Ush2a*^*−/−*^ (*n* = 10 afferents from 7 mice) and *Ush2a*^*+/+*^ mice (*n* = 11 afferents from 6 mice; *P* > 0.05, two-way ANOVA with Bonferroni post-hoc analysis). **e**, Firing activity of Aβ-fiber SAMs in response to the static phases of ramp and hold stimuli did not differ between *Ush2a*^*−/−*^ (*n* = 12 afferents from 6 mice) and *Ush2a*^*+/+*^ mice (*n* = 12 afferents from 5 mice; *P* > 0.05, two-way ANOVA with Bonferroni post-hoc analysis). **f,** Similarly, Aδ-fiber mechanoreceptor responses to static ramp and hold stimuli did not differ (*Ush2a^−/−^ n* = 22 afferents from 6 mice, *Ush2a^+/+^ n* = 14 afferents from 6 mice; *P* > 0.05, two-way ANOVA with Bonferroni post-hoc analysis). **g,** C-fiber nociceptors in response to the static phases of ramp and hold stimuli also did not differ between *Ush2a*^*−/−*^ (*n* = 19 afferents from 6 mice) and *Ush2a*^*+/+*^ mice (*n* = 14 afferents from 8 mice; *P* > 0.05, two-way ANOVA with Bonferroni post-hoc analysis). **h,** Behavioral responses of mice to brushing the hindpaw glabrous skin did not differ between *Ush2a*^*+/+*^ (*n* = 8) and *Ush2a*^*−/−*^ mice (*n* = 12; *P* > 0.05, *t*-test). **i**, Behavioural nocifensive withdrawal thresholds to von Frey hair (vFh) stimulation of the hindpaw glabrous skin did not differ between *Ush2a*^*+/+*^ (*n* = 8) and *Ush2a*^*−/−*^mice (*n* = 12; *P* > 0.05, *t*-test). Data = mean± SEM.

Genetic ablation of *Ush2a* in mice is associated with morphological disruption of the stereocilia, the sites of mechanotransduction in cochlear hair cells^12^. We thus investigated whether genetic ablation of *Ush2a* causes structural impairments in mechanoreceptors or in the end-organs in the skin that they innervate. Ablation of the mouse *Ush2a* gene was not associated with any loss of sensory axons as determined anatomically, nor any changes in myelin thickness (Extended data Fig. 8a-c). More importantly, Meissner corpuscles and hair follicles showed no obvious morphological alterations in *Ush2a*^*−/−*^ mice compared to controls (Extended data Fig 8d-i). We quantified the number of sensory axons per Meissner corpuscle and found it was unchanged in the hind limb glabrous skin (Extended data Fig 8f), despite clear reductions in the mechanosensitivity of RAMs (Extended data Fig 4 f-h). On average hair follicle bulb thickness or Awl/zig-zag and Guard hairs were quantitively indistinguishable between *Ush2a*^*−/−*^ and *Ush2a*^*+/+*^ mice (Extended data Fig 8g-i). We also quantified the number, size and branching of terminal Schwann cells around Awl/zig-zag and Guard hairs from both genotypes and found no quantitative differences (Extended data Fig 8j,k,l). Thus the continued presence of the USH2A protein in mature sensory end-organs and the lack of impact on morphology following genetic ablation suggested that USH2A does not play a role in maintaining the anatomical integrity of the mechanoreceptor end-organ.

Since we only observed functional deficits in forepaw RAMs innervating Meissner corpuscles, we reasoned that the perceptual deficit displayed by *Ush2a*^*−/−*^ mice could primarily be due to sparse input from these neurons during the detection task. We could not record mechanoreceptor activity during the behavioral task, so we instead compared temporal features of mechanoreceptor firing to the time course of the behavioral response (Fig 6). Peristimulus histograms of first lick latencies showed that the many correct responses of both *Ush2a*^*+/+*^ and *Ush2a*^*−/−*^ occurred within 500ms of stimulus onset (Fig 6a,b,e,f; Extended data Fig 9a). Median first lick latencies were consistently longer in *Ush2a*^*−/−*^ compared to mice Usher Peristimulus histograms of mechanoreceptor firing showed that in *Ush2a*^*−/−*^ compared to *Ush2a*^*+/+*^ mice (Extended data Fig 9a), but these differences were not statistically different. RAM activity was almost completely absent with a 12mN 5Hz vibration stimulus and greatly attenuated at 24mN compared to controls (Fig. 6b,d,g,h). In contrast, the firing rates of SAMs measured to the same stimuli were almost indistinguishable between genotypes (Fig. 6c,d,g,h). This data suggests that firing in RAMs is critical for the animals to perform the task and that firing in SAMs, although present, may not provide sufficient information for the animal to reliably perform the task. Interestingly, in the case of a 5Hz stimulus the animal experienced not more than 3 full vibration cycles (1 cycle at 5Hz lasts 200ms) before in the majority of cases acting correctly to receive a water reward (Fig 6 a,c). We thus analysed the temporal features of mechanoreceptor firing in the 600ms epoch prior to the animal’s decision. We found as expected that RAMs show strong phase locking to the sinusoidal stimulus (Extended data Fig 9b-i), similar to primate and human mechanoreceptors^31–33^. We compared firing in mechanoreceptors from control and *Ush2a*^*−/−*^ mice to estimate the minimal amount of information necessary for the mouse to make a decision. It was clear that strong phase locking in a population of RAMs to the first sinusoids is correlated with the animal’s ability to correctly detect the stimulus. Vibration stimuli of high frequency (e.g. 50 Hz) evoked more spikes in RAMs in the period before stimulus detection when applied with sufficient intensity (Extended data Fig 9b-i). We conclude that the reduction in RAM sensitivity to vibration frequencies from 5-50 Hz may be sufficient to explain the perceptual deficit in humans and mice following genetic ablation of *USH2A*. It appeared that humans have a more profound perceptual deficit when confronted with higher frequency vibration (125Hz) than did mice (Fig 1c). Human Meissner corpuscle afferents are very sensitive to such high frequencies^25^ and it may be that there is a species difference regarding the range of vibration frequency to which humans and mice are most sensitive. For technical reasons we were unable to use sinusoids with frequencies greater than 50Hz to stimulate mouse mechanoreceptors. However, it was clear from our analysis that mouse RAMs in the forepaw are in fact particularly well tuned to lower frequency sinusoids (Fig 2g).

**Fig. 6.**
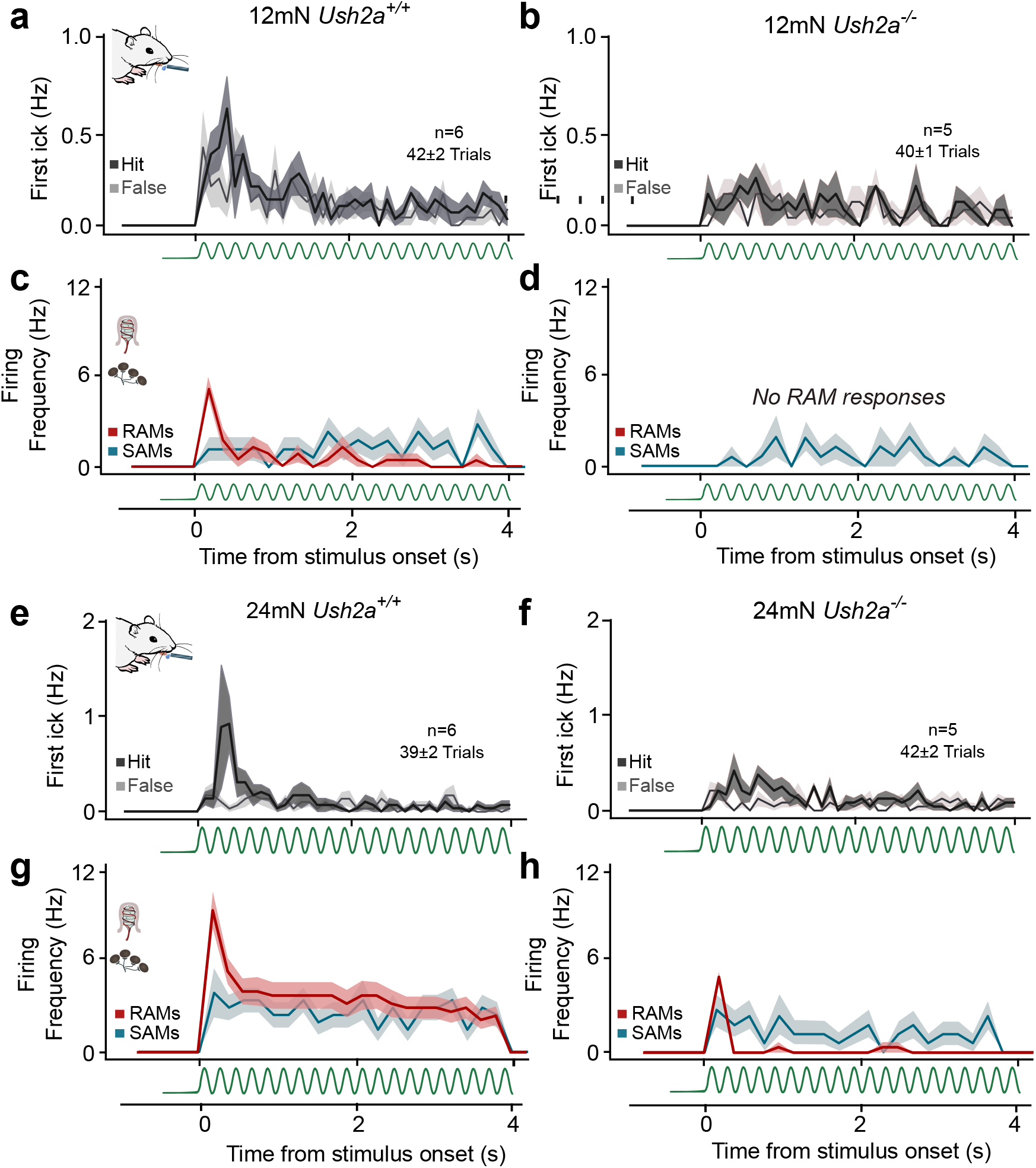
Impaired forepaw RAM mechanosensitivity underlies 5Hz vibration perceptual deficits in *Ush2a*^*−/−*^ mice. **a,** Average PSTHs of first lick responses from all *Ush2a*^*+/+*^ mice during all trials during 12mN (42±2 trials in 6 mice) to 5Hz vibration stimuli. Hit responses above false alarm lick rates indicate stimulus detection within the first 3 sinusoids. **b,** Average PSTHs of first lick responses from all *Ush2a*^*−/−*^ mice during all trials during12mN vibration stimuli (40±1 trials in 5 mice). Note that mice did not correctly report the stimulus, indicated by a lack of difference between first lick hit and false lick rates. **c,** PSTHs of individual forepaw RAMs and SAMs recorded *ex vivo* in *Ush2a*^*+/+*^ mice in response to the same 12mN 5Hz stimuli. (RAM n = 5/11 neurons; SAM n = 5/10 neurons). **d,** PSTHs of RAMs and SAMs recorded in *Ush2a*^*−/−*^ mice. No RAMs responded to 12mN vibration (*n* = 0/10), but SAMs showed vibration sensitivity in the *Ush2a*^*−/−*^ mice at 12mN (*n* = 3/11). **e,** Average PSTHs of first lick responses from all *Ush2a*^*+/+*^ mice during all trials during 24mN 5Hz vibration (39±2 trials in 6 mice). Note the high first lick rates from the second sinusoidal wave. **f,** PSTHs of first lick responses from all *Ush2a*^*−/−*^ mice during all trials during 24mN 5Hz vibration (42±2 trials in 5 mice). *Ush2a*^*−/−*^ mice reported 24mN vibration, as indicated by higher hit vs false rats at the start of the stimulus. **g,** PSTHs of RAMs and SAMs recorded in *Ush2a*^*+/+*^ mice. RAMs responded robustly to 24mN vibration, especially in response to the first sinusoidal wave. (*n* = 9/10). SAMs also responded robustly to 24mN vibration (*n* = 3/11). **h,** PSTHs of RAMs and SAMs recorded in *Ush2a*^*−/−*^ mice. RAMs only fired a single action potential at the start of the 24mN vibration stimulus (*n* = 6/10). SAMs showed normal vibration sensitivity in the *Ush2a*^*−/−*^ mice at 24mN (right; *n* = 10/11). All data = mean± SD.

Here we identified the tether-like protein USH2A as being localized exclusively to end-organ cells where its presence is required to maintain normal vibration sensitivity. It is increasingly apparent that specialized skin cells, e.g. Merkel cells^34,35^ and sub-epidermal Schwann cells^18^, can themselves sense mechanical stimuli that they pass on to sensory endings to mediate touch and pain sensation^18,35,36^. The precise role of classical end-organ structures like the Meissner corpuscle has, however, remained enigmatic. Here we provide molecular insight into the function of such end-organs as the expression of the USH2A protein in cells within the corpuscle adjacent to the mechanoreceptor endings is necessary for these neurons to detect low amplitude vibration. Mechanosensitive currents in mouse and fly sensory neurons are critically dependent on extracellular tethers for normal gating^37–39^. It is thus conceivable that the large extracellular domain of USH2A may underlie a physical link between mechanosensitive channels and associated proteins in the sensory ending^16,40^ to facilitate vibration transduction. It is unlikely that the USH2A protein is identical to a 100 nm long extracellular protein filament that we previously showed to be necessary for mechanosensitivity in all types of mechanoreceptors^37,38^. The USH2A protein does not contain a furin cleavage site, is not expressed by sensory neurons and is restricted to end-organs that are only associated with RAMs. Our data does not prove that the USH2A protein forms part of an intercellular tether complex. With confocal microscopy we could detect the USH2A protein throughout the terminal Schwann cell cytoplasm and at sites of contact with the sensory ending within the Meissner corpuscle (Fig 2a-c). Also, in contrast to auditory hair cells^41^, the exact location of the sites of mechanotransduction in the comparatively vast multicellular end-organ complex of touch receptors is unknown. We observed a highly specific loss of vibration sensitivity associated with loss of USH2A in the absence of any overt anatomical disruption of the end-organ. Loss of USH2A may lead to changes in the stiffness of elements within the Meissner corpuscle this could be due to changes in cellular stiffness or in the stiffness of physical connections between the sensory ending and USH2A-positive Schwann cells within the end-organ. Changes in stiffness could decrease the efficiency with which mechanical forces are propagated to mechanotransduction channels. We conclude that USH2A is likely a critical component of the end-organ that could have a similar function to or interact with intercellular tethers that function like the hair cell tip link^42^.

## Materials and Methods

### Human participants

Experiments in Spain were reviewed and approved by the Ethics Committee of the University Hospital La Fe Valencia and adhered to the tenets of the Declaration of Helsinki and further reviews. Experiments in Berlin were approved by the ethic committee of Charité-Universita◻tsmedizin, Berlin (EA4/012/05). Two cohorts were studied: one consisting of 65 healthy adult volunteers; and one group of 16 Usher Syndrome (type II) patients carrying different truncation mutations in *USH2A*. Data from three patients were omitted from the study due to two patients with prior diagnosis of Diabetes, and one patient with carpal tunnel syndrome, which could affect psychophysical thresholds. Study participants were tested with a battery consisting of nine psychophysical tests applied to the skin. All participants received written and oral information for taking part in the study. The following participant information was documented: age, gender, handedness, hearing devices (hearing aids, cochlear implants), severity of deaf and blindness symptoms, and comorbidities.

### Human quantitative sensory testing

Psychophysical testing was performed according to the standardised testing protocol of the quantitative sensory testing (QST) of the German Research Network on neuropathic pain (DFNS) ^43^. Vibrotactile perceptual thresholds at different frequencies were determined using a two-choice assay ^3,4^. Mechanical vibration stimuli were delivered to the skin between the nail bed and the first joint of the little finger with a piezo-actuator (Physik Instrumente PI) controlled by Powerlab 4/30 and Labchart 7.1 software. The probe was made of glass and had a 5mm diameter flat, circular contact area with a brass counterweight to ensure the same degree of holding force was applied in every subject. The little finger of the dominant hand was tested over two frequencies (10Hz and 125Hz; 1.8s duration). Subjects signalled the detection of a vibratory stimulus during one of two time windows by pressing a button. The protocol consisted of an up-down design that increased or decreased the amplitude of the stimulus (between 18nm to 45μm), depending on the success rate during the task. Eight up/down amplitude reversals (a total of ~40-80 individual trials per session) around the perceptual threshold were performed to create a mean perceptual threshold for each patient. The amplitude of the vibration stimuli driven by the piezo device was calibrated by measuring the displacement amplitude of the probe under a microscope to step voltage increments. A tactile acuity test was also used to test the limits of spatial resolution of the fingertip (median innervation), using a two-interval forced-choice tactile grating orientation test using a Tactile Acuity Cube which consisted of 6 sides with a gratings of different widths (0.75, 1.25, 1.75, 3.0, 4.5 and 6.0mm)^3–5^. Subjects were blindfolded the cube was placed in the vertical or horizontal orientation against their fingertip. A two-down and one-up procedure was employed with 10-15 grating size turning points. Thresholds (71% correct response) were calculated as median from the last 10 of 15 turning points ^4^. Mechanical detection thresholds were determined with a standardised set of 23 von Frey hair filaments (vFhs; Optihair3-Set, Marsock Nervetest, Germany) with forces between 0.25-265mN. Von Frey hairs were applied in an ascending order on the dorsal aspect of the hand (radial innervation) for 1 second until the subject perceived a tactile sensation. The order was then reversed until the point were the subject did not perceive a tactile sensation. The mean force of five reversals was taken as threshold. Mechanical pain thresholds were tested with a set of 7 weighted pinprick stimulator (MRC systems, Germany) with tips of 0.25mm and forces between 8-512mN. A simple staircase method (1 up and 1 down rule) was performed where patients were asked whether the stimulus was perceived as sharpness that causes the feeling of pricking pain. Stimuli were applied to the dorsal aspect of the hand after vFh detection tasks. The threshold was calculated from the mean of five reversals. Thermal warm and cool perception thresholds and hot and cold pain thresholds were tested using a TSA II Thermosensory analyser (MEDOC, Israel). The thermode had an area of 9cm^2^, cut off temperatures between 0-50°C and a temperature change rate of 1°C/s. The thermode was placed on the volar aspect of the mid-forearm region (medial antebrachial innervation). Three consecutive trials were performed for each thermal test in the following order: cold detection, warm detection, and cold pain and heat pain thresholds. Study participants were asked to indicate at what point they start to experience cooling, warming, cold and heat pain. Thresholds were the average temperature of the three trials in each test.

### Mice

All experiments were approved by the Berlin animal ethics committee (Landesamt für Gesundheit und Soziales (LAGeSo), Berlin, Germany) and carried out in accordance with European animal welfare law. Male and female *Ush2a*^*−/−*^ mice and wild type litter mates (*Ush2a*^*+/+*^) from the CBA/CaJ background between 10-30 weeks of age were used ^12^. All mice were maintained on a 12h light/ 12h dark cycle.

### Mouse tissue anatomy and Immunohistochemistry

For skin immunohistochemistry, mice were euthanized by CO_2_ inhalation for 2-4min followed by cervical dislocation, and paw skin tissue was dissected and the hypodermis, ligaments and attached muscle tissue was removed. Skin samples were stretched out using insect pins and fixed for 45 mins in 4% paraformaldehyde (PFA). For gelatine vibratome sections, tissue samples were placed into embedding moulds (Polysciences, T-8), filled with warm gelatine (20%, dissolved in 0.1M PBS), and post-fixed in 4% PFA overnight at 4ºC before 120μm sections were cut using a vibratome (Leica, VT100S). For whole mount staining, skin samples were stretched for 2 hours in PFA, then post-fixed at 4°C for 24h in 20% DMSO and 80% methanol. After cutting or post-fixation, tissue samples were washed thrice in PBS (0.1M) and incubated for 72h at 4°C in blocking solution and primary antibodies. Samples were then washed thrice in PBS before 24h incubation (4°C) with secondary antibodies diluted in blocking solution. Tissue was then again washed thrice, and processed for tissue clearing using 2,2’-Thiodiethanol (TDE, Sigma Aldrich). Sections were placed in increasing concentrations of TDE every 2h, from 10%, 25%, 50% to 97%, in which samples were stored and mounted onto slides and coverslipped. Polyclonal antibodies targeting Ush2A were generated in the rabbit by Eurogentec^®^ (Liege, Belgium). Four antibodies were created that bind to different binding epitopes on the N- and C-termini of Ush2A (N-terminus antibody 1: CSPLYNDKPFRSGDNV and antibody 2: C+SWEKPAENFTRGEII; C-terminus antibody 1: C+ADTRLPRSGTPMSIR and antibody 2: CIRERPPLVPLQKRMT), and were purified from anti-sera before use. All four antibodies were used together (1:200) for staining protocols in sequential stainings with Chicken anti-neurofilament 200 (NF200, Millipore, 1:1000), Rabbit anti-S100 (Dako, 1:1000) or Guinea Pig anti-CK20 (Origene Tech, Germany, 1:200). Secondary antibodies used were anti-rabbit Alexa Fluor 488, anti-chicken Alexa Fluor 647, anti-rabbit Alexa Fluor 647 and anti-Guinea Pig Alexa Fluor 647 (Invitrogen, 1:800). Tiled Z-stack images were obtained using a confocal microscope (LSM700 Carl-Zeiss) using Zen2009 software. NF200+ and S100+ Nerve fiber innervating Meissner corpuscles and hair follicles were visualized and counted using Fiji/ImageJ. For electron microscopy images of the sciatic nerve, animals were perfused and the sciatic nerve was dissected and fixed in 4% PFA and 2.5% glutaraldehyde and contrasted with osmium tetroxide before embedding in Technovit 7100 resin (Heraeus Kulzer, Wehrheim, Germany). Ultra-thin sections captured at 5600x magnification. Myelinated nerve fibers and unmyelinated fibers were counted and measured using iTEM software.

### Mouse skin-nerve preparation and sensory afferent recordings

Cutaneous sensory fiber recordings were performed using the *ex vivo* skin nerve preparation. Mice were euthanized by CO_2_ inhalation for 2-4min followed by cervical dislocation. Three different preparations were performed in separate experiments using different paw regions: the saphenous nerve innervating the hairy hindpaw skin ^44^; the Tibial nerve innervating the glabrous hindpaw skin; and the medial and ulnar nerves innervating the forepaw glabrous skin ^20^. In all preparations, the hairy skin of the limb was shaved and the skin and nerve was dissected free and transferred to the recording chamber where muscle, bone and tendon tissues were removed from the skin to improve recording quality. The recording chamber was perfused with a 32°C synthetic interstitial fluid (SIF buffer): 123mM NaCl, 3.5mM KCl, 0.7mM MgSO_4_, 1.7mM NaH_2_PO_4_, 2.0mM CaCl_2_, 9.5 mM sodium gluconate, 5.5mM glucose, 7.5mM sucrose and 10mM HEPES (pH7.4). The skin was pinned out and stretched, such that the outside of the skin could be stimulated using stimulator probes. The peripheral nerve was fed through to an adjacent chamber in mineral oil, where fine filaments were teased from the nerve and placed on a silver wire recording electrode.

The receptive fields of individual mechanoreceptors were identified by mechanically probing the surface of the skin with a blunt glass rod or blunt forceps. Analog output from a Neurolog amplifier were filtered and digitized using the Powerlab 4/30 system and Labchart 7.1 software (ADinstruments). Spike-histogram extension for Labchart 7.1 was used to sort spikes of individual units. Electrical stimuli (1Hz, square pulses of 50-500ms) were delivered to single-unit receptive fields to measure conduction velocity and enable classification as C-fibers (velocity <1.2ms^−1^), A-delta fibers (1.2-10ms^−1^) or A-beta fibers (>10ms^−1^). Mechanical stimulation of the receptive fields of neurons were performed using a piezo actuator (Physik Instrumente, P-841.60) and a double-ended Nanomotor (Kleindiek Nanotechnik, MM-NM3108) connected to a force measurement device (Kleindiek Nanotechnik, PL-FMS-LS). Calibrated force measurements were acquired simultaneously using the the Powerlab system and Labchart software during the experiment.

As different fiber types have different stimulus tuning properties, different mechanical stimuli protocols were used based on the unit type. Low threshold Aβ-fibers and Aδ-fiber D-hairs were stimulated with 3 vibration stimuli (5Hz, 25Hz and 50Hz) with increasing amplitude over 6 steps (from ~6-65mN; 20 cycles per step), and a dynamic stimulus sequence with four ramp and hold waveform with varying probe deflection velocity (3s duration; 0.075mms^−1^, 0.15mms^−1^, 0.45mms^−1^ and 1.5mms^−1^; average amplitude 100mN). Aβ-fiber slowly-adapting mechanoreceptors (SAMs) and rapidly-adapting mechanoreceptors (RAMs) were classified by the presence or absence of firing during the static phase of a ramp and hold stimulus, respectively as previously described^20,21^. Single-units were additionally stimulated with a series of five static mechanical stimuli with ramp and hold waveforms of increasing amplitude (3s duration; ranging from ~10-260mN). High threshold Aδ-fibers and C-fibers were also stimulated using the five ramp and hold stimuli with increasing amplitudes.

### Single-unit recordings from Pacinian afferents

To assess the mechanosensitivity of Pacinian corpuscles located around the fibula, the skin and all muscles were removed from the lower leg and the interosseous nerve was dissected free from the interosseous membrane. Subsequently, the fibula and tibia, which in mice are fused along their distal half, were removed from the animal together with the interosseous nerve and were transferred to a custom-made organ bath chamber where they were mounted with a custom-made mini-vice. The entire dissection procedure was performed in ice-cold dissection buffer which contained 108 mM NMDG, 20 mM HEPES, 3.5 mM KCl, 10 mM MgSO4, 26 mM NaHCO3, 1.7 mM NaH2PO4, 9.5 mM Na-gluconate, 5.5 mM Glucose, 18.5 mM sucrose and 0.5 mM CaCl2 (adjusted to pH 7.4 with NaOH). After a recovery period of 10 min, the preparation was perfused with oxygenated recording buffer (35°C) which consisted of 108 mM NaCl, 3.5 mM KCl, 0.7 mM MgSO4, 26 mM NaHCO3, 1.7 mM NaH2PO4, 9.5 mM Na-gluconate, 5.5 mM Glucose, 7.5 mM sucrose and 1.5 mM CaCl2 (adjusted to pH 7.4 with NaOH) and the proximal end of the interosseous nerve was transferred to an oil-filled recording chamber by threading it through a small hole (~1 mm diameter) that connected the organ bath chamber with the adjacent recording chamber. After removal of the perineurium, single filaments were dissected free and attached to the recording electrode for action potential recording. Recordings were made with the Neurolog system (NL100AK, NL104A, Digitimer Ltd, UK) and a PowerLab 4SP controlled by LabChart 7.1 (AD Instruments). To determine the mechanical activation threshold and the frequency tuning of Pacinian afferents, a series of sinusoidal mechanical stimuli (duration 1s; linearly increasing amplitude dAmp/dt = 30μm/s; maximum amplitude 30μm) with increasing frequencies (40-480 Hz) were applied to the distal end of the fibula with a metal rod (tip diameter 1 mm) that was attached to a piezo actuator (P-840.2, Physik Instrumente GmbH, Germany).

### Mouse vibration perception learning task

Firstly, for implantation of the head restraint, mice were anesthetized with isoflurane (1.5-2% in O_2_) and injected subcutaneously with Metamizol (200 mg per kg of body weight). A light metal support was implanted onto the skull with glue (UHU dent) and dental cement (Paladur). Temperature of mice was monitored with a rectal probe and kept at 37°C using a heating pad. Mice were then placed in their home cage with Metamizol (200 mg/ml) in the drinking water. Implanted mice were habituated to head-restraint 3-5 days after surgery. Mice were gradually habituated in the behavioral setup to head fixation and paw tethering using a medical (cloth) tape. Next, one day following the beginning of water restriction mice underwent two pairing sessions on consecutive days.

During pairing sessions (30-45 minute duration), water rewards were given from a water spout paired to presentation of the vibration stimulus (3 second duration, 5Hz, 60mN) to the glabrous skin of the forepaw to build an association between stimulus and reward. The vibration stimulus was given to the restrained paw via a custom made 2mm^2^ cushioned glass rod attached to a piezo actuator (PL127.11 PICMA from Physik Instrumente). A voltage-force relationship was calibrated by using a force measurement system (Dual-Mode Lever Arm system 300-C Aurora Scientific) to measure the applied sinusoidal force profile applied by the piezo device after each experiment.

Following pairing, daily training sessions began where mice received a small water reward (4-7 μl) from the spout when they correctly licked during a timeout upon start of the stimulus (3.5 seconds). Catch trials, where no stimulus was presented and licks were counted as false alarms, were interleaved as 50% of the total trials.Trials were not cued by any external event or stimulus as previously described^11^ and were delivered at randomized time intervals between 3 and 30 s. If mice licked during a 2s window before the stimulus onset, a 3-30s delay was imposed to promote stimulus-water reward association. One training session consisted of around 60 trials (30x stimulus + 30x catch). Hits and false alarm rates were compared to assess performance during the training sessions. To test that mice were licking in response to vibration stimulus of the forepaw, one session was included at the end of training where the stimulating device was moved 3-5mm below the paw so no skin contact was made. In subsequent experimental days after mice had successfully learned the task, vibration stimuli were given with different amplitudes and frequencies. For 5Hz vibration, 48mN, 36mN, 24mN, 12mN and 6mN forces were used to stimulate the forepaw, with two different amplitudes interleaved with catch trials during each day. For example, 48mN, 24mN stimuli and catch trials were interleaved during one session which consisted of around 100 trials (33x 48mN stimulus + 33x 24mN stimulus + 34x catch). 6mN stimuli were tested over two sessions. For 25Hz and 125Hz vibration, 60mN and 12mN forces of the same frequency were interleaved with catch trails in the same manner over two testing sessions.

### Mouse paired pulse inhibition behavioral assay

The startle response of mice was assessed using the Startle Response system (SR-LAB, San Diego Instruments). Mice were habituated to the apparatus for 30mins each day for 3 days before testing. Briefly, mouse startle responses were measured during a total of 48 pulse-alone trials^45^ (40ms duration, 129dB) and prepulse + pulse trials (20ms prepulse of 69dB, 73bD or 81dB). Prepulse trials were 200ms before pulse trials. The percentage paired pulse inhibition (% PPI) for each prepulse intensity was calculated as %PPI = 100 × ([(pulse-alone) − (prepulse + pulse)]/pulse-alone.

### Mouse von Frey hair and brush behavioral assays

Mice were habituated to the testing environment for 30mins each day for 3 days before testing. On experimental days, mice were placed in individual Plexiglas cubicles on an elevated mesh platform. For the brush assay ^46^, the left hindpaw was stroked from heel to toe with a 2mm head paint brush 10 times at random intervals. The percentage of withdrawals was calculated. On subsequent testing days, mechanical nocifensive withdrawal thresholds were measured using von Frey hairs. Briefly, the left hindpaw plantar surfaces were stimulated with von Frey hair filaments and the reflex threshold was determined by using the up-down method ^47^. Depending on the response of the animal, higher or lower force von Frey hairs were applied to the hindpaw to establish a series of 6-9 positive or negative reflex withdrawals. This pattern of responses was then converted such that data is expressed as log of the mean of the 50% reflex withdrawal threshold ^47^.

### Data analysis of human psychophysics data

Data were tested for normality, and differences in psychophysical performance for each test between control participants and patients with Usher syndrome were compared using unpaired t-test or Mann-Whitney tests. Z-transformations were used to compare single individual data profiles with the healthy control mean and standard deviation. The Z-score was calculated using excel spreadsheets based on the formula Z=(patient score – group mean /standard deviation of the group mean). Z-values above 0 indicate gain of function, meaning the subject is more sensitive to the tested stimuli compared to healthy controls. Individual Z-scores lying outside the 95% confidence interval (i.e. Z-score <1.96 or >1.96 standard deviation) of the healthy control dataset can be identified. Performance in each test was tested in pairwise comparisons using the Mann-Whitney U-test and we considered p<0.01 to be statistically significant.

### Data analysis of mouse vibration behaviour

Licks were recorded with a sensor at the tip of the water reward spout. In stimulus trials, a hit was counted when there was a lick within the window of opportunity (3.5 seconds) after the start of the stimulus. Behavioral data was collected using custom-written routines in Lab View at 1 kHz sampling rate, and custom-written Python scripts were used for analysis. During catch trials, a false alarm took place when there was a lick during an equally long window of opportunity. To assess whether mice successfully learnt the detection task, hit rates were compared to false alarm rates within the same training session using repeated measures 2-way ANOVAs with Bonferroni post hoc testing.

To quantify performance in the detection tasks, we used d’ (sensitivity index) instead of the percentage of correct trials in order to take into account bias in the licking criteria ^48^. To calculate d’, the following formula was used: d’ = z(h) – z(fa), where z(h) and z(fa) are the normal inverse of the cumulative distribution function of the hit and false alarm rates, respectively. To avoid infinity d’ values, when all trials were reported (rate = 1) or none of them was (rate = 0), the rates were replaced by (1-1/2N) or (1/2N), respectively, where N is the number of trials the stimulus was presented ^49^. The Z-scores for hit and false alarm rates were calculated with OpenOffice Calc (Apache Software Foundation) using the function NORMINV. We have only performed statistical tests on data where at least one of the genotypes shows a d’ value above 1.0 indicating that these mice reported the stimulus. Behavioral data was collected using custom-written routines in Lab View at 1 kHz sampling rate, and custom-written Python scripts were used for analysis. The custom codes and scripts are available upon request.

### Data analysis of skin-nerve preparation recordings

Cutaneous forepaw and hindpaw units were categorized based on their conduction velocity and responses to mechanical stimuli. Mechanical thresholds of units were calculated as the temperature or mechanical amplitude required to elicit the first action potential. All statistical analyses were performed with GraphPad Prism 6.0 and Python. Statistical tests include two-way repeated measures ANOVA with Bonferroni’s post hoc test, Student t-test, Mann Whitney test and Wilcoxon matched pairs test. Kolmogorov-Smirnov test was used to assess normality of the data. Asterisks in figures indicate statistical significance: *p<0.05, **p<0.01, ***p<0.001.

### Data analysis of mouse nerve electron microscopy

For analysis of mouse Sciatic nerve electron micrographs, numbers of myelinated and unmyelinated fibers were counted in 12 sections per nerve. The total numbers of fibers per nerve were then extrapolated from the cross sectional area of each nerve ^50^.

## Acknowledgements

We thank Maria Braunschweig and Heike Thränhadt for technical assistance, Dr. Tiansen Li for providing us with *Ush2a*^*−/−*^ mice, Dr. Bettina Purfürst for electron microscopy, Melani Navarro for assistance with patient recruitment and Annapoorani Udhayachandran for advice. This study was funded by grants from the Deutsche Forschungsgemeinshaft (GRL SFB665-B6; JFAP, SFB 1315; SGL SFB1158-A01) and grants from European Research Council grant to (G.R.L, ERC 789128: JFAP, ERC-2015-CoG-682422;). Additional funding from FIS PI16/00539 from the Institute of Health Carlos III (ISCIII, Spanish Ministry of Science and Innovation) to JM. We thank Ulrich Müller for critical comments on the MS.

## Author Contributions

FS, VB, RM and GRL conceived the human and mouse studies. FS carried out mouse skin electrophysiology and behaviour experiments with guidance from GRL and JFAP. FJT and SGL established, performed and analysed data from Pacinian afferent recordings. FS, BM and TD carried out immunohistochemical and anatomical experiments. VB RM and JK recruited the control cohort, collected and analysed the QST data of the control cohort. Psychophysical experiments with Spanish Usher patients were carried out by VB, FS, RM, GGG, JK and JOA. FS and GRL wrote the manuscript with input from all authors.

## Data Availability

Raw data from this manuscript is available upon request.

## Author Information

Reprints and permissions information are available at www.nature.com/reprints.

Correspondence and requests for materials should be addressed to

---

**Extended Data Fig 1. Mouse vibration perceptual training task.**

**a,** Hit and False alarm lick rates for *Ush2a*^*+/+*^ and **b,** *Ush2a*^*−/−*^ mice during behavioral vibration detection training. *Ush2a*^*+/+*^ and *Ush2a*^*−/−*^ mice learned to report 5Hz 60mN forepaw vibration after 4 days (Hit vs False lick rates within each genotype: \*\*\**P* < 0.001; two-way ANOVA with Bonferroni post-hoc analysis). Light colour lines show data from individual mice, bold lines show population mean.

**c,** Perceptual sensitivity (d’) measurements show how perceptual sensitivity increases during training in both *Ush2a*^*+/+*^ and *Ush2a*^*−/−*^ mice.

**d,** Perceptual thresholds of *Ush2a*^*+/+*^mice to 5Hz stimuli. *Ush2a*^*+/+*^ mice reported 5Hz stimuli ≥12mN, as shown by significantly higher Hit vs False Alarm lick rates ≥12mN (Hit vs False: \*\*\**P* < 0.001; Two-way repeated measures ANOVA with Bonferroni post-hoc analysis).

**e,** *Ush2a*^*−/−*^ mice reported 5Hz vibration stimuli ≥24mN (Hit vs False: \*\*\**P* < 0.001; two-way ANOVA with Bonferroni post-hoc analysis).

**f,** Mean first lick latencies during hit trials of *Ush2a*^*−/−*^ mice were significantly longer at 36mN 5Hz vibration (\**P* < 0.05; Two-way ANOVA with Bonferroni post-hoc analysis).

**g,** Hit and False licking rates of mice during 25Hz vibration detection task in *Ush2a*^*+/+*^ and *Ush2a*^*−/−*^ mice. Dots show data from individual mice, horizontal lines show mean.

**h,** Hit and False licking rates of mice during 100Hz vibration detection task in *Ush2a*^*+/+*^ and *Ush2a*^*−/−*^ mice.

**i,** Mice trained on the vibration detection task did not respond to possible sounds generated by the vibration stimulus when the Piezo device was moved 5mm below the forepaw (*P* > 0.05; paired t-tests). Data = mean ± S.E.M. n = 11 mice. Filled dot shows hit rates, open dot shows false alarm lick rates.

**Extended Data Fig 2. Ush2a^−/−^ mice have a mild hearing deficit in a paired-pulse inhibition behavioural task.**

**a,** Response amplitudes (in arbitrary units) of *Ush2a*^*+/+*^ (n=10) and *Ush2a*^*−/−*^ (n=9) mice to 20-40ms auditory tones of different amplitudes did not differ (*P* > 0.05; Two-way ANOVA with Bonferroni post-hoc analysis). **b,** Lower amplitude tones reduced the startle response when given 200ms before the startling 129dB tone in both *Ush2a*^*+/+*^ and *Ush2a*^*−/−*^ mice, however the amplitude of the response was significantly lower in *Ush2a*^*−/−*^ mice compared to *Ush2a*^*+/+*^ mice at 73dB and 81bD prepulse tones (\*\**P* < 0.001; two-way ANOVA with Bonferroni post-hoc analysis). Data = mean± SEM

**Extended Data Fig 3. USH2A immunolabelling in cutaneous Meissner corpuscles**

**a,** USH2A (green) and NF200 (red) immunolabelling of dorsal root ganglia neurons. **a’** USH2A channel only. **a’’** NF200 channel only. A’’’ RNA scope probe for *Ush2a* mRNA shows no labelling in the DRG (green) a second probe for NF200 mRNA (red) shows large sensory neurons labeled. The *Ush2a* RNA labels Meissner corpuscle cells in the skin (green). **b** NF200 (red), and S100 (cyan) immunolabelling of Meissner corpuscles in the sectioned paw glabrous skin. **b’** USH2A channel. **b’’** NF200 channel. **b’’’** S100. **c,** USH2A (green) and NF200 (red) immunolabelling of deep nerve branches in wholemounted paw skin show an absence of USH2A in nerve bundles. **c’** USH2A channel only. **c’’** NF200 channel. **d,** USH2A (green) and NF200 (red) immunolabelling of Meissner corpuscles in the sectioned paw glabrous skin from an *Ush2a*^*−/−*^ mouse show an absence of USH2A labelling. **d’** USH2A channel only. **d’’** NF200 channel only.

**Extended Data Fig 4. *Ex vivo* forepaw and hindpaw skin nerve preparation recordings from cutaneous RAM Aβ-fibers**

**a,** The percentages of forepaw RAMs that respond to different amplitudes of 5Hz (top) and 50Hz (bottom) vibration stimuli in *Ush2a*^*+/+*^ (*n* = 11 afferents from 5 mice) and *Ush2a*^*−/−*^ mice (*n* = 10 afferents from 4 mice). **b,** *Ush2a*^*−/−*^ forepaw RAMs are significantly less responsive to 10Hz and **c,**50Hz vibration compared to RAMs in *Ush2a*^*+/+*^ mice (\*\**P* < 0.01, *** *P* < 0.001; two-way ANOVA with Bonferroni post-hoc analysis). **d,** Phase-locking (average number of spikes per sinusoid) decreased as vibration amplitude decreased in *Ush2a*^*+/+*^ forepaw RAMs. **e.** Phase-locking to vibration stimuli (5-50 Hz) was generally lower in *Ush2a*^*−/−*^ forepaw RAMs. **f,** Hindpaw glabrous Aβ-fiber RAMs recorded from *Ush2a*^*−/−*^ mice (*n* = 14 afferents from 4 mice) were significantly less responsive to 5Hz (left: *,**P < 0.05, 0.01, two-way ANOVA with Bonferroni post-hoc analysis), **g,** but not 50Hz (right: *P* > 0.05; Two-way ANOVA with Bonferroni post-hoc analysis) vibration than *Ush2a*^*+/+*^ mice *(n* = 13 afferents from 4 mice). **h,** Hindpaw glabrous Aβ-fiber RAMs responses to ramp stimuli of differing velocities were significantly lower in *Ush2a*^*−/−*^ mice (*P < 0.05, two-way ANOVA with Bonferroni post-hoc analysis). **i,** Forepaw Aβ-fiber RAMs *(n* = 11 neurons from 5 mice). were significantly more sensitive to 5Hz and**, j,**50Hz vibration than hindpaw RAMs in *Ush2a*^*+/+*^ mice *(n* = 13 afferents from 4 mice: *,**,***P < 0.05, 0.01, 0.001, two-way ANOVA with Bonferroni post-hoc analysis). **k,** Forepaw and hindpaw Aβ-fiber SAM responses to 5Hz vibration did not statistically differ (*P* > 0.05; Two-way ANOVA with Bonferroni post-hoc analysis). Data = mean± SEM.

**Extended Data Fig 5. USH2A immunolabelling in the mouse skin**

**a,** Wholemount USH2A (green) and S100 immunolabelling of a hair follicle with lanceolate endings in the hindpaw hairy skin. **a’** USH2A channel only. **a’’** S100 channel only. Scale bars 25μm. **b,** Wholemount USH2A (green) and S100 (red) immunolabelling of a hair follicle with circumferential (circ) endings showing USH2A expression in the hindpaw hairy skin. **b’** USH2A channel only. **b’’** S100 channel only. Scale bars 25μm. **c,** Wholemount USH2A (green) and S100 (red) immunolabelling of a hair follicle with circumferential endings that is USH2A-negative in the hindpaw hairy skin. **c’** USH2A channel only. **c’’** NF200 channel only. Scale bars 25μm. **d,** Wholemount USH2A (green) and CK20 (red: marker of Merkel cells) immunolabelling of Guard hair in the back hairy skin with surrounding Merkel cell complex. Note the lack of overlap between USH2A and CK20 immunolabelling. **d’** USH2A channel only. **d’’** S100 channel only. Scale bars 50μm. **e,** USH2A (green) and S100 (red) immunolabelling of D-hairs in sectioned glabrous hindpaw skin, showing absence of USH2A staining in D-hairs. **e’** USH2A channel only. **e’’** S100 channel only. Scale bars 50μm.

**Extended Data Fig 6. Aβ-fibre SAMs and A**δ**-fibre D-hairs have normal mechanosensitivity in *Ush2a*^*−/−*^ mice.**

**a,** Example traces of single SAMs recorded from *Ush2a*^*+/+*^ (top) and *Ush2a*^*−/−*^ (bottom) mice firing in response to the 5Hz vibration protocol. **b,** SAMs recorded from *Ush2a*^*+/+*^ (*n*=10 afferents from 4 mice) and *Ush2a*^*−/−*^ (*n*=11 afferents from 4 mice) mice show comparable sensitivity to 25Hz and **c,**50Hz vibration (*P* > 0.05; Two-way ANOVA with Bonferroni post-hoc analysis. **d,** Aδ-fibre D-hairs recorded from the hairy hindpaw skin and glabrous hindpaw skin in *Ush2a*^*+/+*^ (*n*=9 afferents from 6 mice) and *Ush2a*^*−/−*^ (*n*=11 afferents from 6 mice) mice showed comparable sensitivity to 25Hz and **e,**50Hz vibration (*P* > 0.05; Two-way ANOVA with Bonferroni post-hoc analysis). **f,** D-hair vibration frequency tuning was not significantly different in *Ush2a*^*+/+*^ and *Ush2a*^*−/−*^ mice (*P* > 0.05; Two-way ANOVA with Bonferroni post-hoc analysis) Data = mean± SEM.

**Extended Data Fig 7. Warm and cool detection by forepaw thermoreceptors is not impaired in *Ush2a*^*−/−*^ mice.**

**a,** The percentages of forepaw thermoreceptive C- and A-fibers classified by their response to different stimuli in *Ush2a*^*+/+*^ and *Ush2a*^*−/−*^ mice (n=48 afferents from 6 mice). C-MH = C-mechanoheat; C-MHC = C-mechanoheatcold; C-MC = C-mechanocold; C-C = C-cold; A-MC = A-mechanocold. **b,** Firing rates of cool-sensitive afferents recorded in *Ush2a*^*+/+*^ and *Ush2a*^*−/−*^ mice during a 32-12°C ramp (temperature change = 1°C/second) was not statistically different (*P* > 0.05; two-way ANOVA with Bonferroni post-hoc analysis). **c,** Similarly, firing rates of heat-sensitive thermoreceptors during a 32-48°C ramp did not differ between *Ush2a*^*+/+*^ and *Ush2a*^*−/−*^ mice. Data = mean ± SEM

**Extended Data Fig 8. Tibial nerve sensory neurons and hindpaw skin touch end-organs in *Ush2a*^*−/−*^ mice show no anatomical defects**

**a,** Example ultrathin electron micrograph (EM) of the tibial nerve from an *Ush2a*^*+/+*^ mouse. Myelinated fibers can be distinguished by the thick grey circumference; Remak bundles of C-fibers lack this myelination. **a’,** Representative electron micrograph of the tibial nerve from an *Ush2a*^*−/−*^ mouse. Scale bars = 2μm. **b,** Mean numbers of A and C-fibers in the tibial nerves of *Ush2a*^*+/+*^ (*n* = 3) and *Ush2a*^*−/−*^ mice (*n* = 3) did not significantly differ (*P* > 0.05; one-way ANOVA with Bonferroni post hoc analysis). Data quantified from EM images. **c,** A-fiber myelin thickness in the tibial nerve did not differ between *Ush2a*^*+/+*^ and *Ush2a*^*−/−*^ mice. **d,** Example S100 (green) and NF200 (red) labelling of Meissner corpuscles in the glabrous hindpaw skin of an *Ush2a*^*+/+*^ mouse and **e,** an *Ush2a*^*−/−*^ mouse demonstrated no obvious morphological deficits. Scale bars = 50μm. **f,** The mean number of fibers that innervate each Meissner corpuscle did not differ between *Ush2a*^*+/+*^ (*n* = 5 mice) and *Ush2a*^*−/−*^ mice (*n* = 5 mice: *P* < 0.05; unpaired *t*-test). **g,** Immunolabelling S100 (green) and NF200 (red) labels hair follicle in the hairy hindpaw skin in an *Ush2a*^*+/+*^ mouse. Scale bar 25μm. **h,** S100 (green) and NF200 (red) labelling hair follicles in the hairy hindpaw skin of an *Ush2a*^*−/−*^ mouse demonstrated no obvious morphological deficits. Scale bar 25μm. Data = mean± SEM. **i,** Average hair bulb thickness of awl/zigzag hairs and Guard hairs did not differ between *Ush2a*^*+/+*^ and *Ush2a*^*−/−*^ mice (unpaired t-tests, P>0.05) Similarly, **j,** the number of terminal Schwann cells per hair follicle, **k,** the average terminal Schwann cell soma size, **l,** and branch lengths did not differ differ between *Ush2a*^*+/+*^ and *Ush2a*^*−/−*^ mice (unpaired t-tests, P>0.05)

**Extended data Fig. 9. Impaired forepaw RAM mechanosensitivity underlies 25Hz vibration perceptual deficits in Ush2a−/− mice**

**a**, Box and whisker plot of median first lick latencies of Ush2a+/+ (n=6) and Ush2a−/− (n=5) mice at 5Hz, 25Hz and 100Hz frequency vibration. **b** PSTHs of first lick responses to 25Hz 12mN vibration stimuli from all behaving *Ush2a*^*+/+*^ and *Ush2a*^*−/−*^ trials. Peak first lick Hit responses above False lick rates indicate first responses of mice 25Hz stimuli. *Ush2a*^*−/−*^ mice did not detect this stimulus **c** Spike timing raster plots of individual forepaw RAMs and SAMs recorded in response to the first 20 cycles of a 25Hz vibration stimulus in the ex vivo forepaw skin nerve preparation of *Ush2a*^*+/+*^ and *Ush2a*^*−/−*^ mice. Each row represents one trial from a single afferent 1-3 trials per recording (left *Ush2a*^*+/+*^ RAM n = 4/11, SAM 2/10 afferents; right *Ush2a*^*−/−*^ RAM n = 1/10, SAM 1/11 afferents). **d**, PSTHs of mean first lick responses to 60mN 25Hz vibration stimuli from all behaving *Ush2a*^*+/+*^ and *Ush2a*^*−/−*^, both genotypes detected this stimulus. **e**, Spike timings of individual RAMs and SAMs recorded in the ex vivo forepaw skin nerve preparation in *Ush2a*^*+/+*^ and *Ush2a*^*−/−*^ mice in response to the first 20 cycles of 60mN 25Hz stimuli (left *Ush2a*^*+/+*^ RAM n = 10/11, SAM n = 10/10; right *Ush2a*^*−/−*^ RAM n = 8/10, SAM n = 10/11). Data = mean ± SD

**Supplementary Movie 1**

Z-projection of the Meissner pictured in Figure 2a.

**Supplementary Movie 2**

Z-axis projections of a single Meissner corpuscle labelled for USH2A (green), S100B (magenta), and NF200 sensory axons (Blue).

**Supplementary Movie 3**

Z-axis projections of a single Meissner corpuscle labelled for USH2A (green), S100B (magenta), and NF200 sensory axons (Blue).

