## Supplementary figures and images for "USH2A is a skin end-organ protein necessary for vibration sensing in mice and humans"

### Extended Data Figure 1

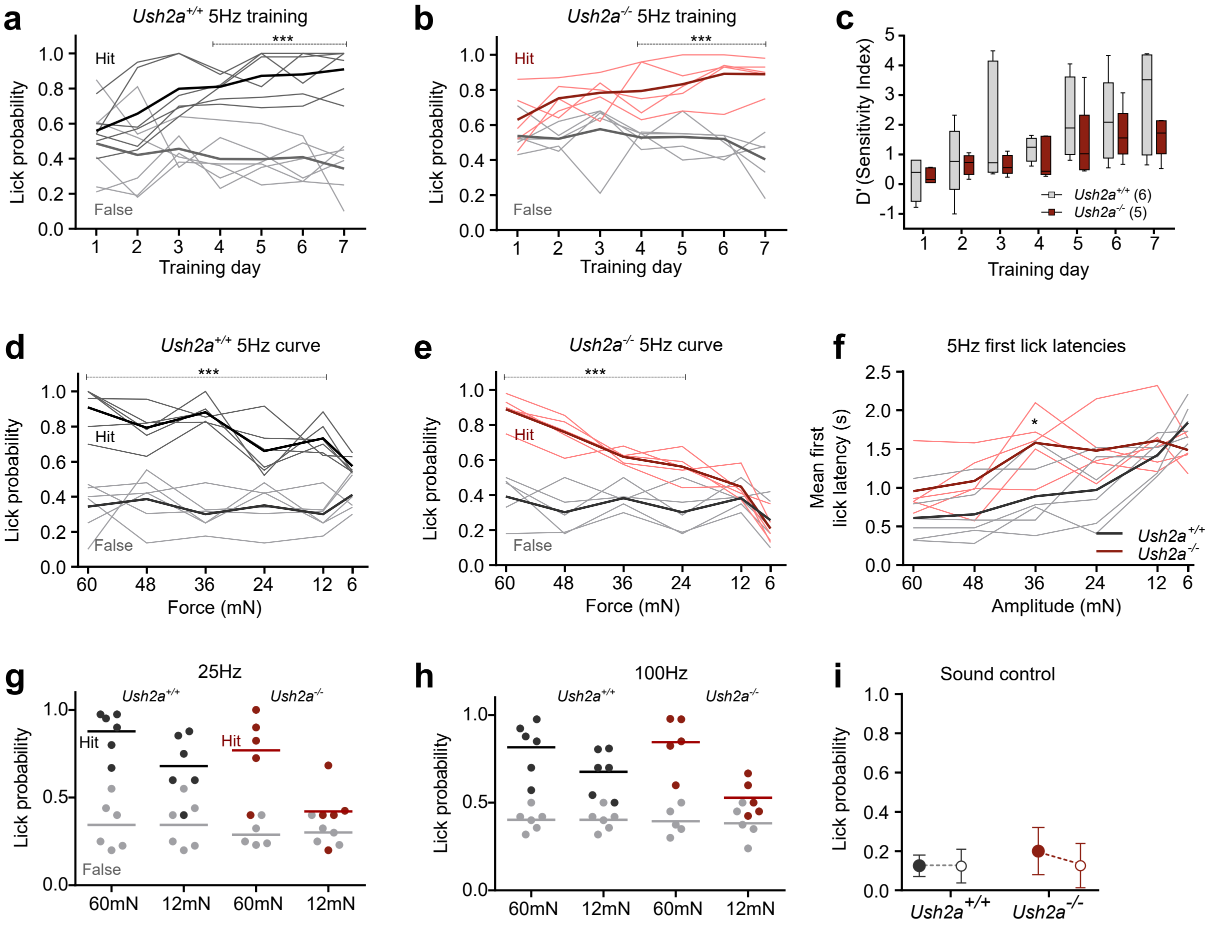

### Extended Data Figure 2

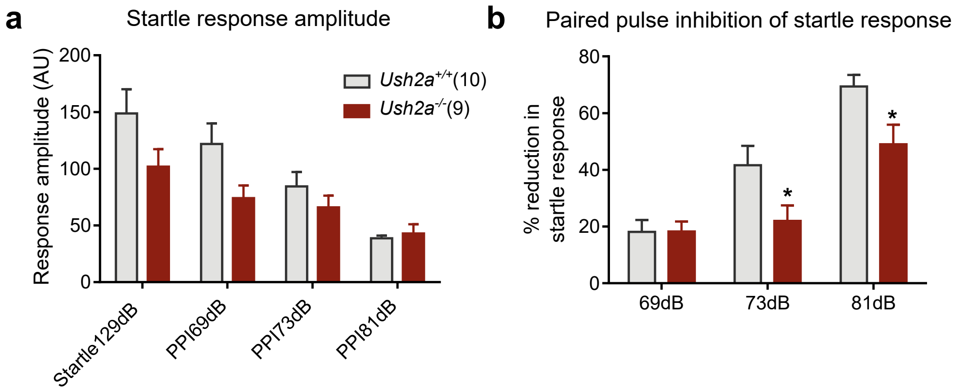

### Extended Data Figure 3

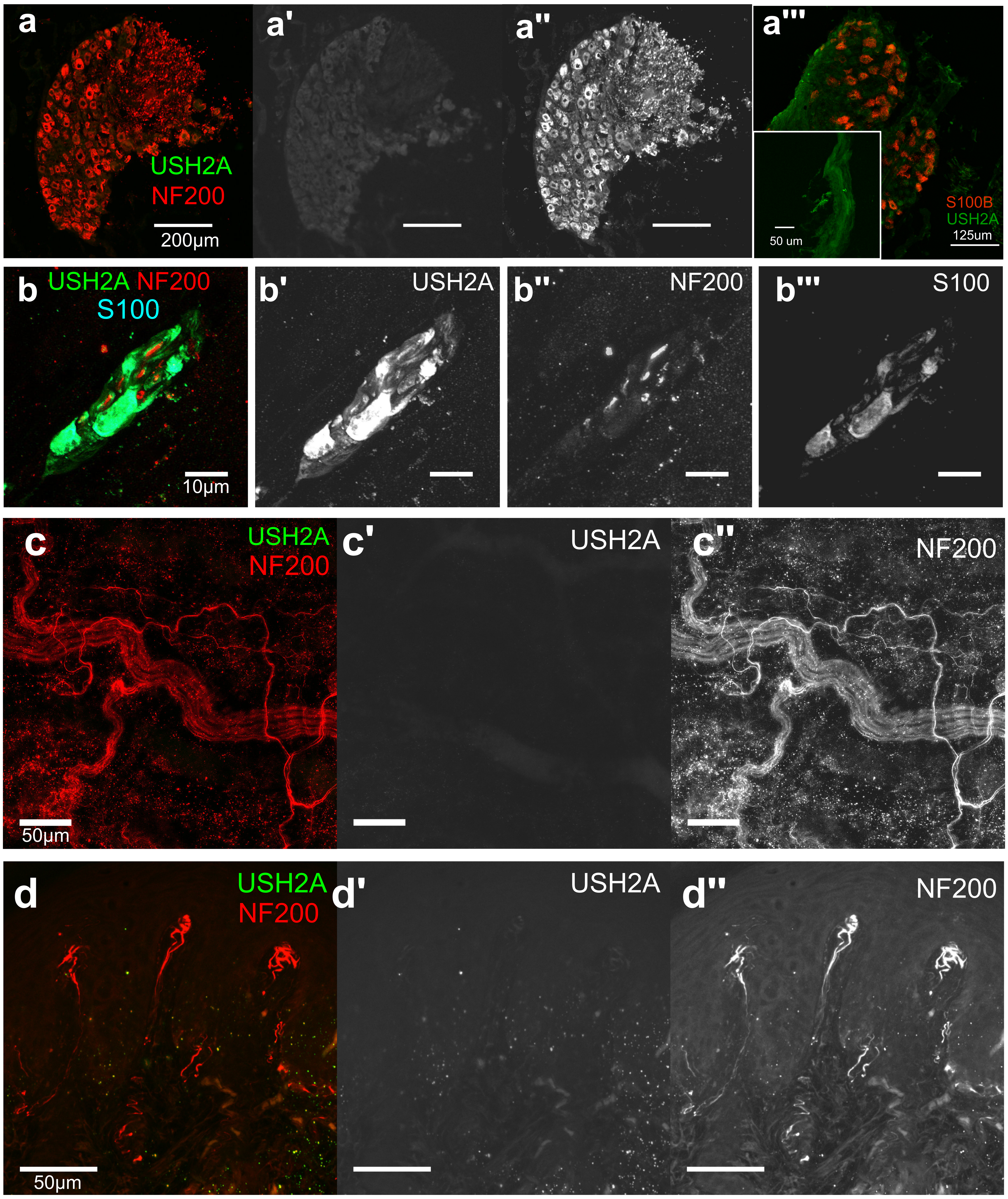

### Extended Data Figure 4

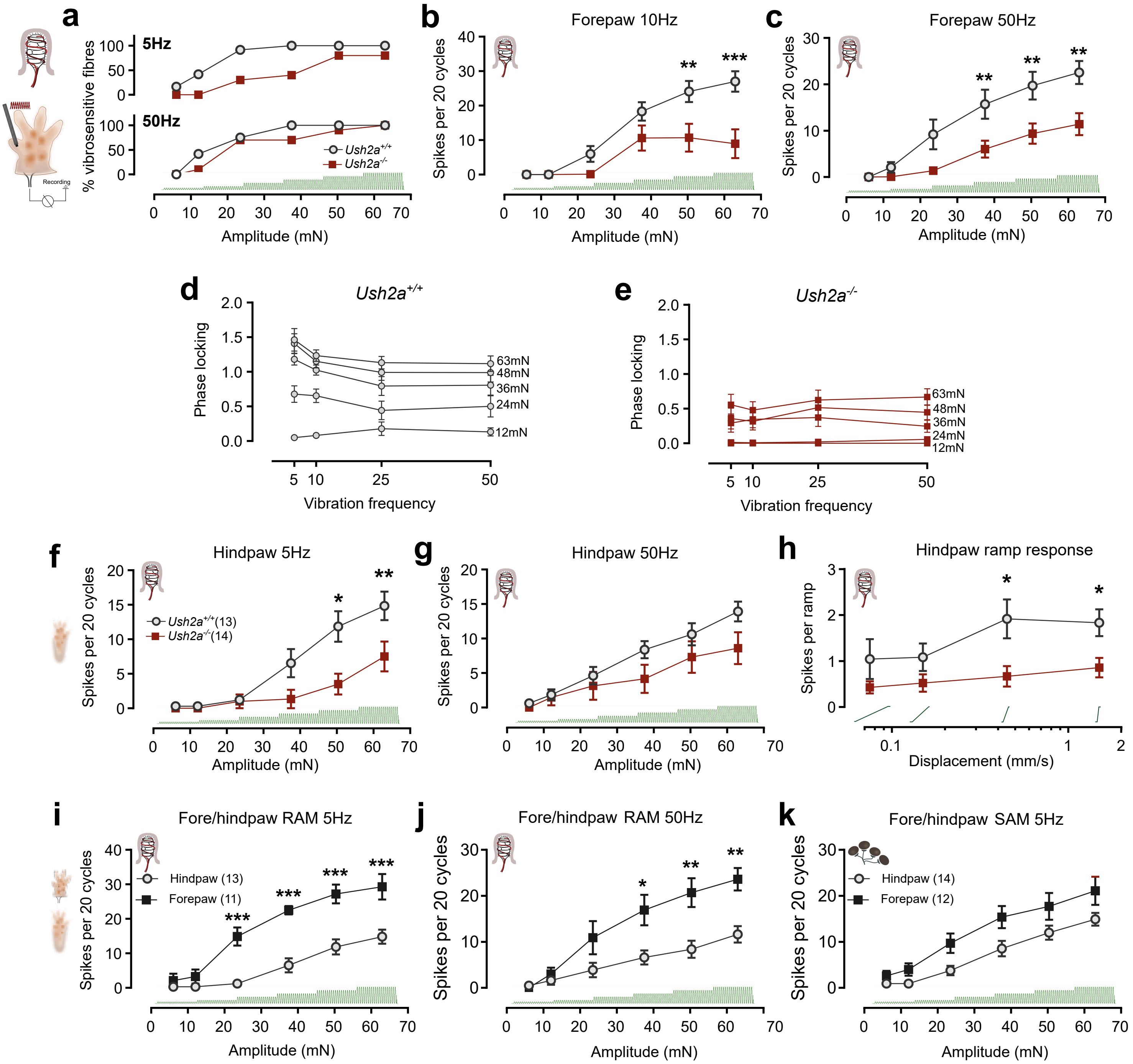

### Extended Data Figure 5

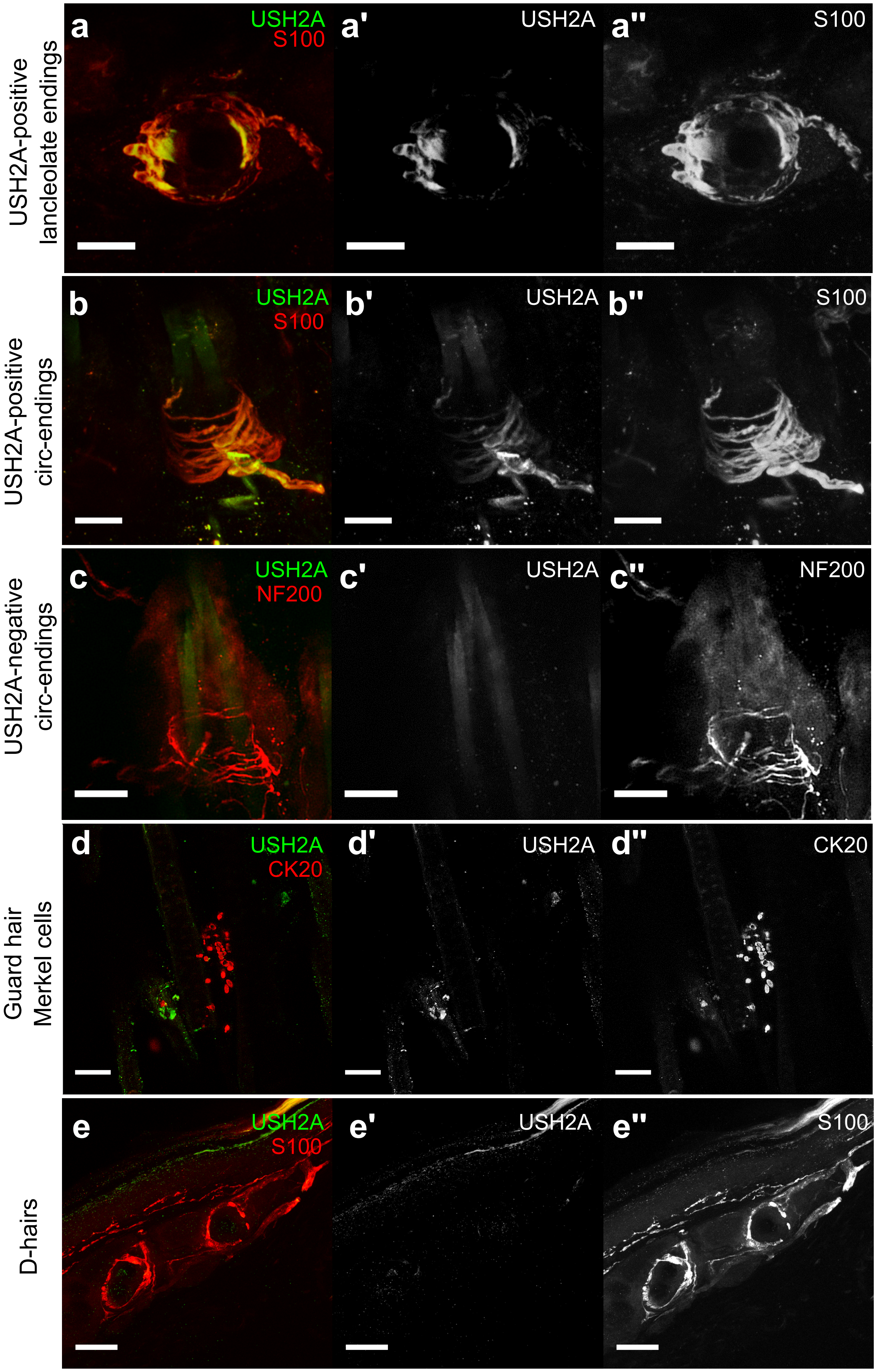

### Extended Data Figure 6

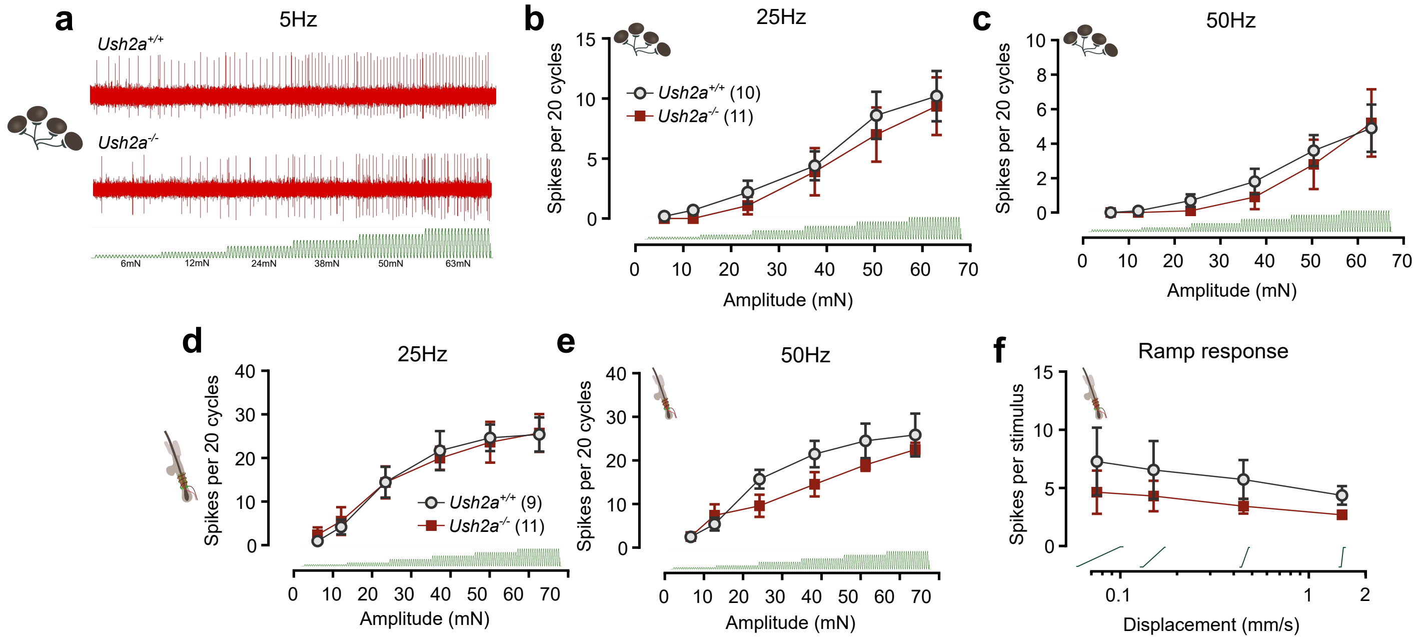

### Extended Data Figure 7

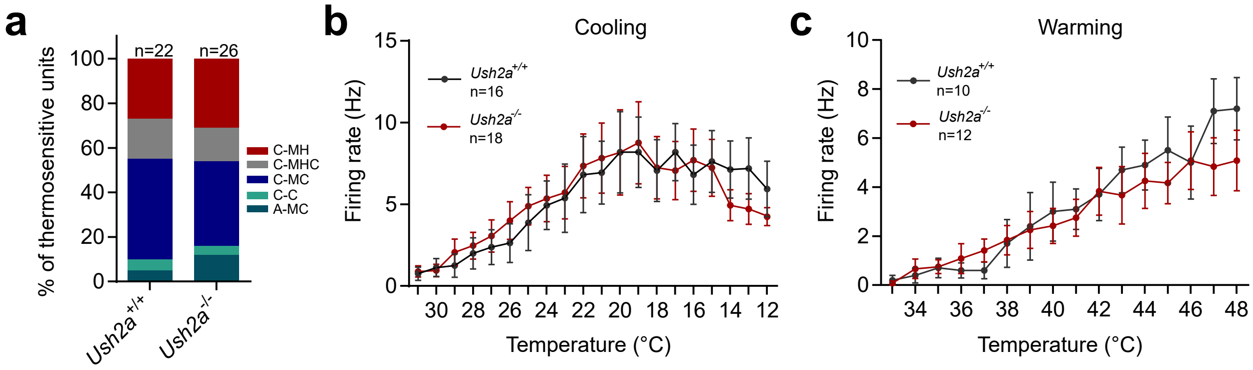

### Extended Data Figure 8

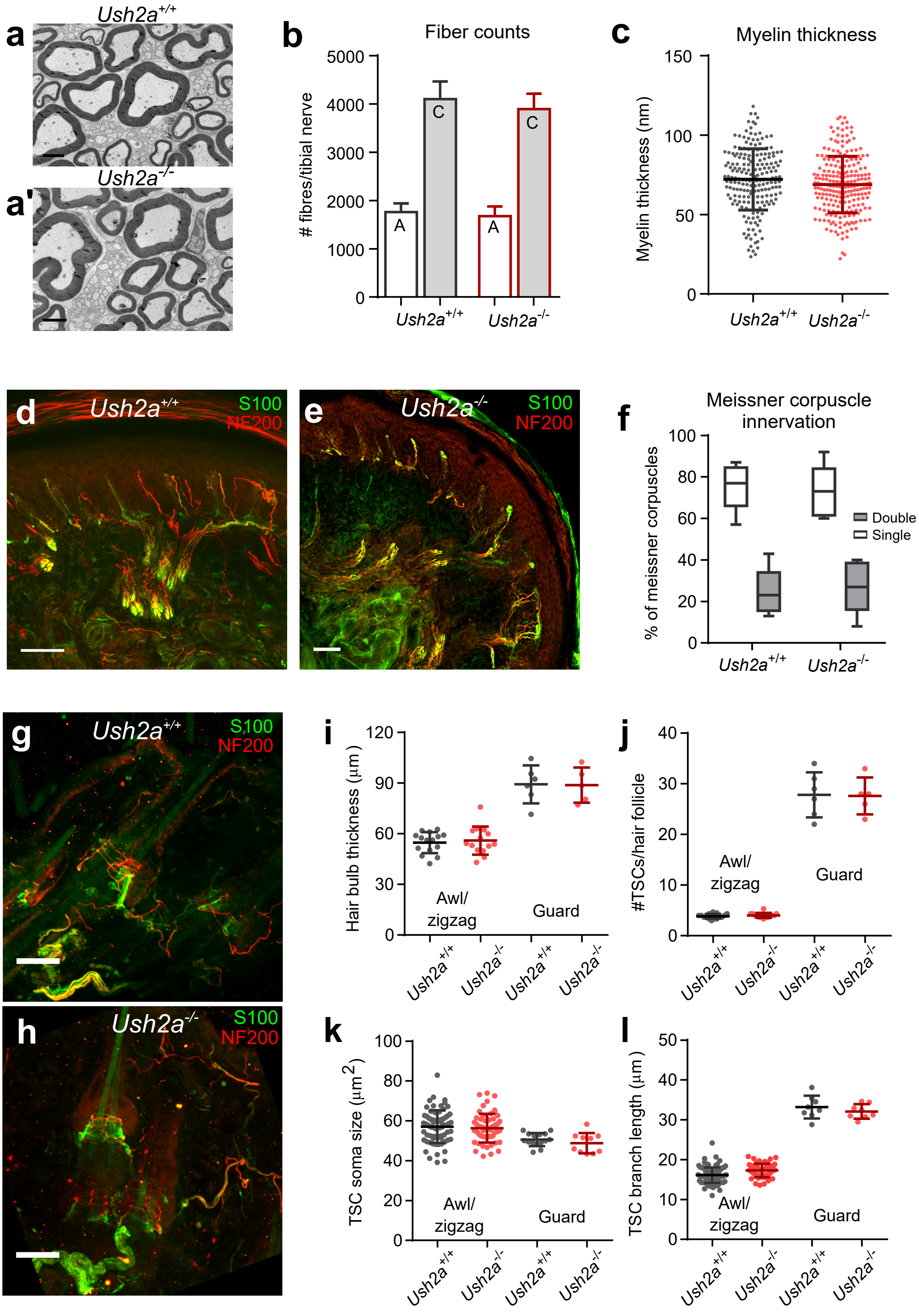

### Extended Data Figure 9

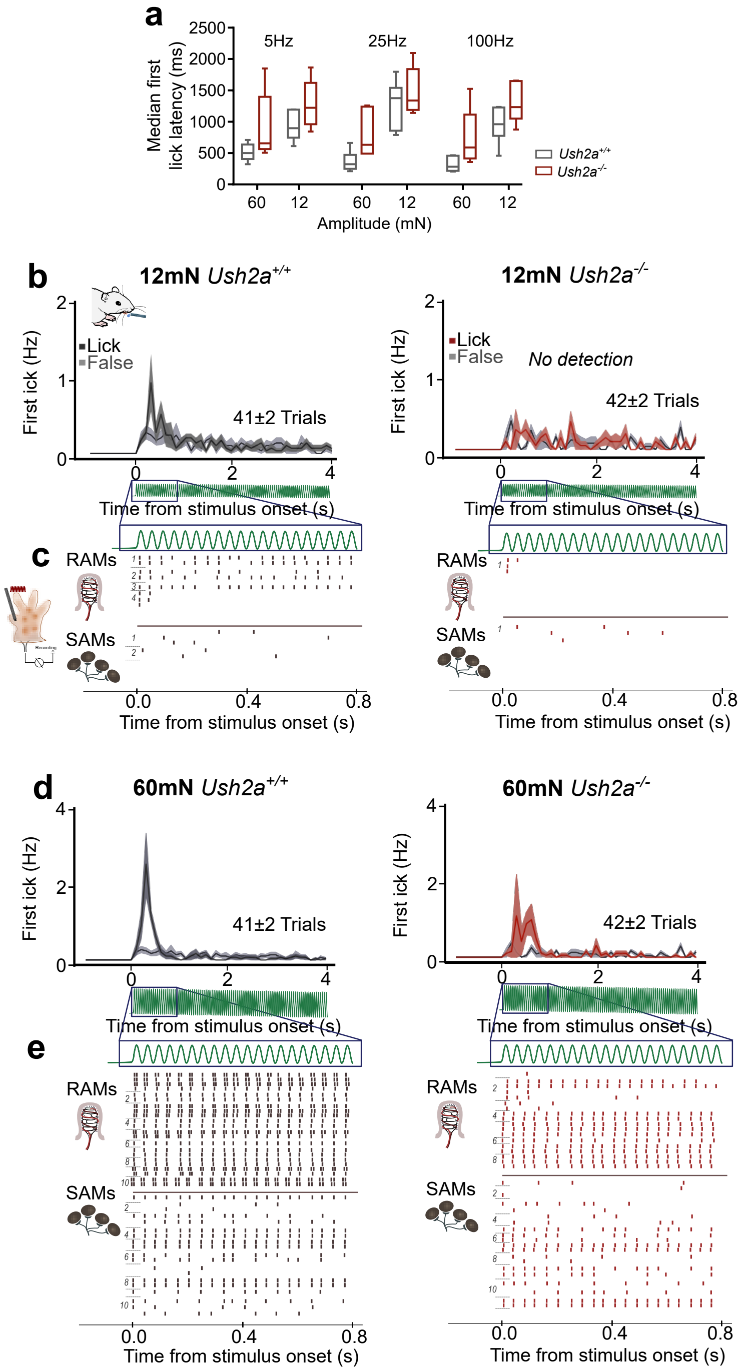

### Supplementary Movie 1

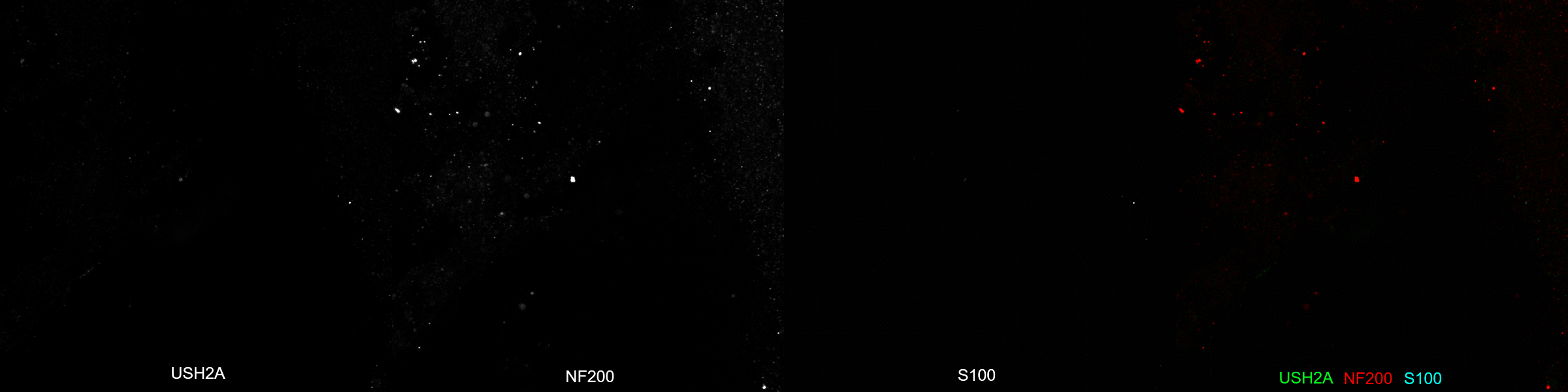

### Supplementary Movie 2

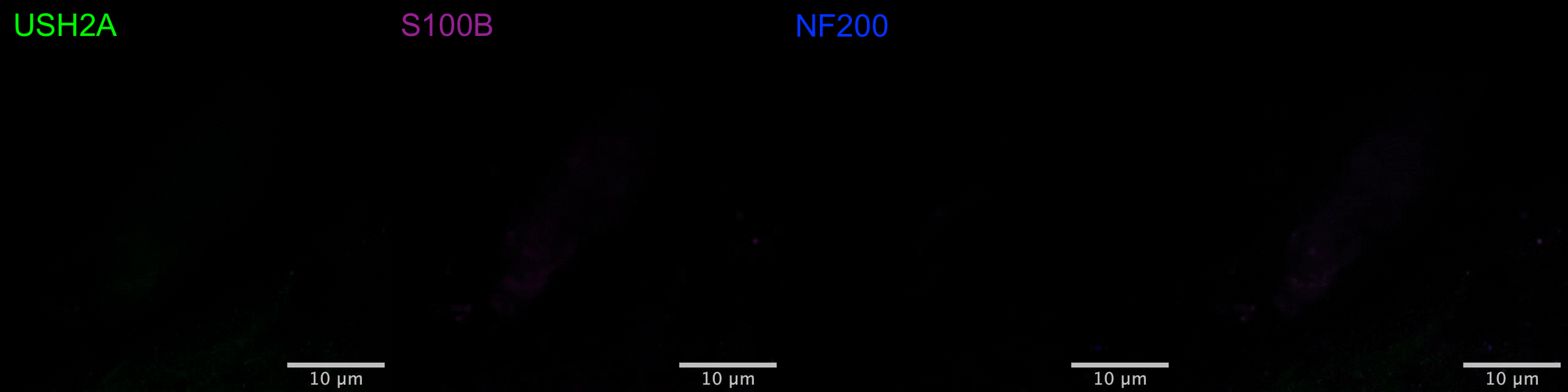

### Supplementary Movie 3

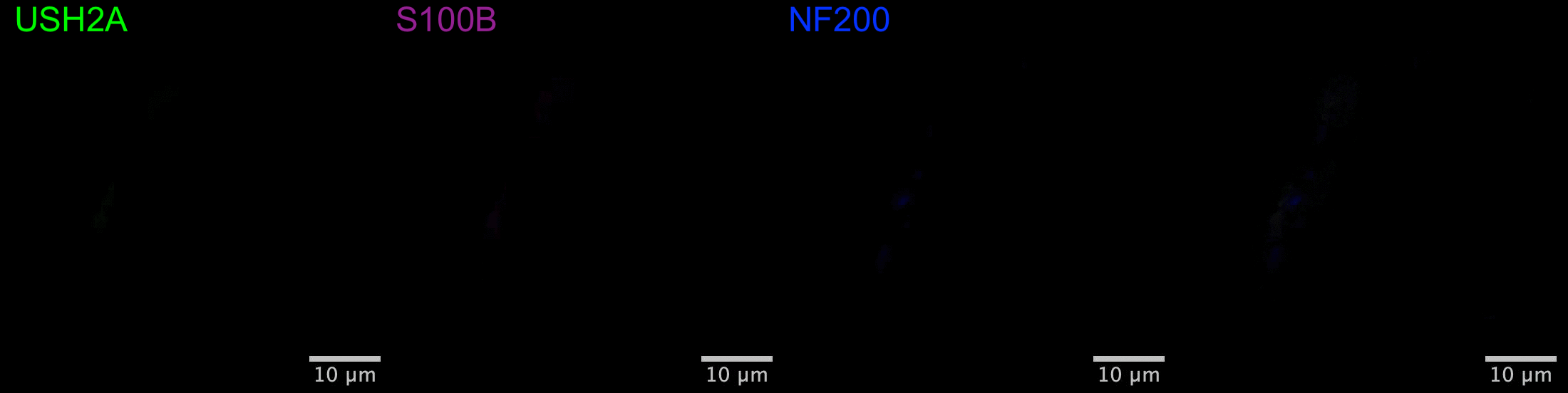
