## Extended data Table 1 for "USH2A is a skin end-organ protein necessary for vibration sensing in mice and humans"

|  |  |  | **Gender distribution** | | **handedness** | |
| --- | --- | --- | --- | --- | --- | --- |
| **Cohort** | **size** | **mean age**  **± SD (year)** | **female** | **male** | **left hand** | **right hand** |
| **control** | N = 65 | 37.75 ±12.92 | 43 (66%) | 22 (34%) | 5 (7.7%) | 60 (92.3%) |
| **patient** | N = 13 | 44.36 ±12.25 | 8 (61.5%) | 5 (38.5%) | 0 | 13 (100%) |

**Supplementary Table 1** Age and sex distribution of patient and control cohort

There was no statistically significant difference in the mean age of the control and patient cohort (Kolmogorov-Smirnov test: P = 0.2). Ten patients out of 13 had retinitis pigmentosa. None of the patients had vestibular symptoms.
