## Extended data Table 2 for "USH2A is a skin end-organ protein necessary for vibration sensing in mice and humans"

|  | | **Usher Syndrome** | | **Control** | | **Mann Whitney tests** |
| --- | --- | --- | --- | --- | --- | --- |
|  |  | **Mean** | **95% CI** | **Mean** | **95% CI** | **Patient vs control** |
| **10Hz vibration** | µm | 6.319 | 4.153-8.486 | 4.085 | 3.505-4.666 | ***P*=0.0026 |
| **125Hz vibration** | nm | 0.947 | 0.438-1.457 | 0.228 | 0.171-0.285 | ****P*<0.0001 |
| **vFh detection** | mN | 2.020 | 1.534-2.506 | 1.386 | 1.145-1.627 | *P*=0.0138 |
| **Pinprick** | mN | 198.3 | 65.93-150.6 | 108.1 | 89.43-126.9 | *P*=0.6784 |
| **Tactile acuity** | mm | 1.346 | 1.211-1.481 | 1.730 | 1.593-1.868 | ***P*=0.0043 |
| **Cool detection** | ΔTºC | 1.616 | 1.603-2.169 | 1.066 | 0.940-1.191 | **P*=0.0088 |
| **Warm detection** | ΔTºC | 1.925 | 1.672-2.179 | 1.877 | 1.734-2.021 | *P*=0.5177 |
| **Cold pain** | ºC | 15.48 | 9.571-21.38 | 10.42 | 8.254-12.58 | *P*=0.0136 |
| **Heat pain** | ºC | 41.62 | 39.18-44.05 | 44.78 | 43.99-45.56 | *P*=0.0514 |

**Supplementary Table 2. Human psychophysical testing data**

Mean values, confidence intervals (CI) and statistical significance of Mann Whitney tests for psychophysical comparisons. vFh = von Frey hair. *P value <0.01, **P value <0.005 ***P value p<0.001
