## Extended data Table 3 for "USH2A is a skin end-organ protein necessary for vibration sensing in mice and humans"

|  |  |  | **VDT**  **10 Hz** | **VDT**  **125 Hz** | **Cooling threshold** |
| --- | --- | --- | --- | --- | --- |
| **Patient** | **USH2A allele 1** | **USH2A allele 2** | **μm** | **μm** | **ΔTC°** |
| 1 | c.2299delG (UV4) | c.2299delG (UV4) | 4.50 | 0.4 | 1.61 |
| 2 | c.2299delG (UV4) | c.2299delG (UV4) | 8 | 2.53 | 1.15 |
| 3 | c.2299delG | c.2299delG | 15.97 | 1.59 | 2.6 |
| 4 | c.2299delG (UV4) | c.11234dupA (UV4) | 28.4 | 0.57 | 1.74 |
| 5 | c.2299delG (UV4) | c.11234dupA (UV4) | 6.36 | 0.32 | 3.94 |
| 6 | c.2299delG (UV4) | DEL EXON 20 (UV4) | 10.08 | 1.13 | 1.57 |
| 7 | c.2299delG (UV4) | p.R303S (UV4) | 3.19 | 0.06 | 1.34 |
| 8 | c.2299delG | c.949C>A | 1.6 | 0.22 | 1.31 |
| 9 | c.4474G>T (p.Glu1492X) (UV4) | c.269A>G (p.Tyr90Cys) (UV3) | 6.36 | 0.28 | 2.61 |
| 10 | c.2431_2432delAA (UV4) | c.2135delC (UV4) | 6.36 | 0.23 | 0.64 |
| 11 | c.8435_8438delCCTA (p.Thr2812Metfs*17) | c.7595-2144A>G | 4.5 | 1.29 | 1.13 |
| 12 | c.5540dupA (p.N1848EfsX20) (UV4) | Del IVS4-IVS9 | 4.5 | 2.53 | 0.9 |
| 13 | c.5540dupA (UV4) | Del IVS4-IVS9 | 5.67 | 0.9 | 0.92 |

**Supplementary Table 3:** Mutation type and psychophysical scores of patients in this study. Most of the mutations were confirmed by sanger sequencing. VDT: vibration detection threshold given in μm**;** cooling threshold given in ΔTC°. To compare with Mean value ± SD of the control cohort: VDT 10 Hz control Mean = 4.1 ± 2.3; VDT 125 Hz control Mean = 0.23 ± 0.23; cooling threshold control Mean = 1.07 ± 0.50.
